# Interfering with a memory without erasing its trace

**DOI:** 10.1101/783654

**Authors:** Gesa Lange, Mario Senden, Alexandra Radermacher, Peter De Weerd

## Abstract

Previous research has shown that performance of a novice skill can be easily interfered with by subsequent training of another skill. We address the open questions whether extensively trained skills show the same vulnerability to interference as novice skills and which memory mechanism regulates interference between expert skills. We developed a recurrent neural network model of V1 able to learn from feedback experienced over the course of a long-term orientation discrimination experiment. After first exposing the model to one discrimination task for 3480 consecutive trials, we assessed how its performance was affected by subsequent training in a second, similar task. Training the second task strongly interfered with the first (highly trained) discrimination skill. The magnitude of interference depended on the relative amounts of training devoted to the different tasks. We used these and other model outcomes as predictions for a perceptual learning experiment in which human participants underwent the same training protocol as our model. Specifically, over the course of three months participants underwent baseline training in one orientation discrimination task for 15 sessions before being trained for 15 sessions on a similar task and finally undergoing another 15 sessions of training on the first task (to assess interference). Across all conditions, the pattern of interference observed empirically closely matched model predictions. According to our model, behavioral interference can be explained by antagonistic changes in neuronal tuning induced by the two tasks. Remarkably, this did not stem from erasing connections due to earlier learning but rather from a reweighting of lateral inhibition.

## 1 Introduction

Expert motor and perceptual skills have been reported to show little decay over periods of months and even years, despite a lack of further practice (Fisk, Hertzog, Lee, Rogers, & Anderson-Garlach, 1994; Hikosaka et al., 2002; Avi Karni & Sagi, 1993; Park, Dijkstra, & Sternad, 2013). Overlearning, reaching automaticity, and fluency during acquisition have been suggested to increase resistance to decay (Dougherty & Johnston, 1996; Farr, 1987; Hagman & Rose, 1983; Healy et al., 1993; Healy, Fendrich, & Proctor, 1990). If true, novice skills may be more vulnerable to interference than expert skills. The instability of novice skills is supported by studies showing the erasure of gains acquired in a single session of training in a first task, when another task is trained in a single session within ∼4h on the same day (Brashers-Krug, Shadmehr, & Bizzi, 1996; Krakauer, Ghilardi, & Ghez, 1999; Shadmehr, Brandt, & Corkin, 1998; Shadmehr & Holcomb, 1997). In these experiments, retention of the first task was impaired on the next day of testing, an effect referred to as behavioral interference. Related time-limited interference has been reported in successive-learning studies involving training over only a few sessions, with interference occurring for time intervals from 0 up to 4h (Seitz et al., 2005; Yotsumoto, Chang, Watanabe, & Sasaki, 2009; J.-Y. Zhang et al., 2008). Caithness et al. (2004), interestingly, showed interference among novice skills for much longer periods. Expert skills, however, typically require extended practice over many more sessions (Ericsson & Lehmann, 1996; Kaufman & Kaufman, 2007; Schoups, Vogels, & Orban, 1995; Schoups, Vogels, Qian, & Orban, 2001; Simon & Chase, 1973; Vogels & Orban, 1985). In view of reports that expert skills resist decay (Fisk et al., 1994; Hikosaka et al., 2002; Avi Karni & Sagi, 1993; Park et al., 2013), the question arises whether highly trained skills are also resistant to interference, perhaps because longer and repeated training as well as consolidation (Wickelgreen, 1972) put expert skills in a ‘permastore’ (Bahrick, 1984).

To the best of our knowledge, no studies tested the stability of expertise by first applying extensive training in one task to then test susceptibility to interference of the acquired skill by extensive training in another task. Here, we trained human participants for 15 daily sessions, spread over about a month, in a visual orientation discrimination task with a given set of orientations (Task A). The participants had to judge whether a Gabor stimulus deviated clockwise or counterclockwise from a (never shown) reference orientation. We then trained the participants for 15 sessions using a different set of orientations (Task B). We tested whether returning to the original set of orientations (Task A) allowed participants to return to the expert level of performance reached after the first training period, and if not, whether further training would restore the originally reached performance level. In addition to the above-described ABA design, we also used a BAB design, as well as a control experiment in which no interference was expected.

To determine the conditions under which interference might occur, we used the finding that extensive training in orientation discrimination until the phase where learning becomes asymptotic, lastingly affects orientation tuning curves in early visual cortex (Raiguel, Vogels, Mysore, & Orban, 2006; Schoups et al., 2001; Yang & Maunsell, 2004). Specifically, as illustrated in figure 1a,b, the flanks of tuning curves overlapping with a trained reference orientation steepen after extensive training in orientation discrimination (Schoups et al., 2001), while the opposite flanks flatten (Teich & Qian, 2003). Steepened flanks permit increased differential activity for stimuli around the trained reference, supporting better discrimination. Accordingly, interference might be induced by additional training with another reference overlapping with the flattened flank, because this can be expected to undo the sharpening that supported the performance increase in the initial training (figure 1b,c). Interference would hence be expected when two tasks are trained to expert-levels that lead to plastic changes in low-level areas including V1, where the two tasks require opposite tuning changes in the same neuronal population.

**Figure 1:**
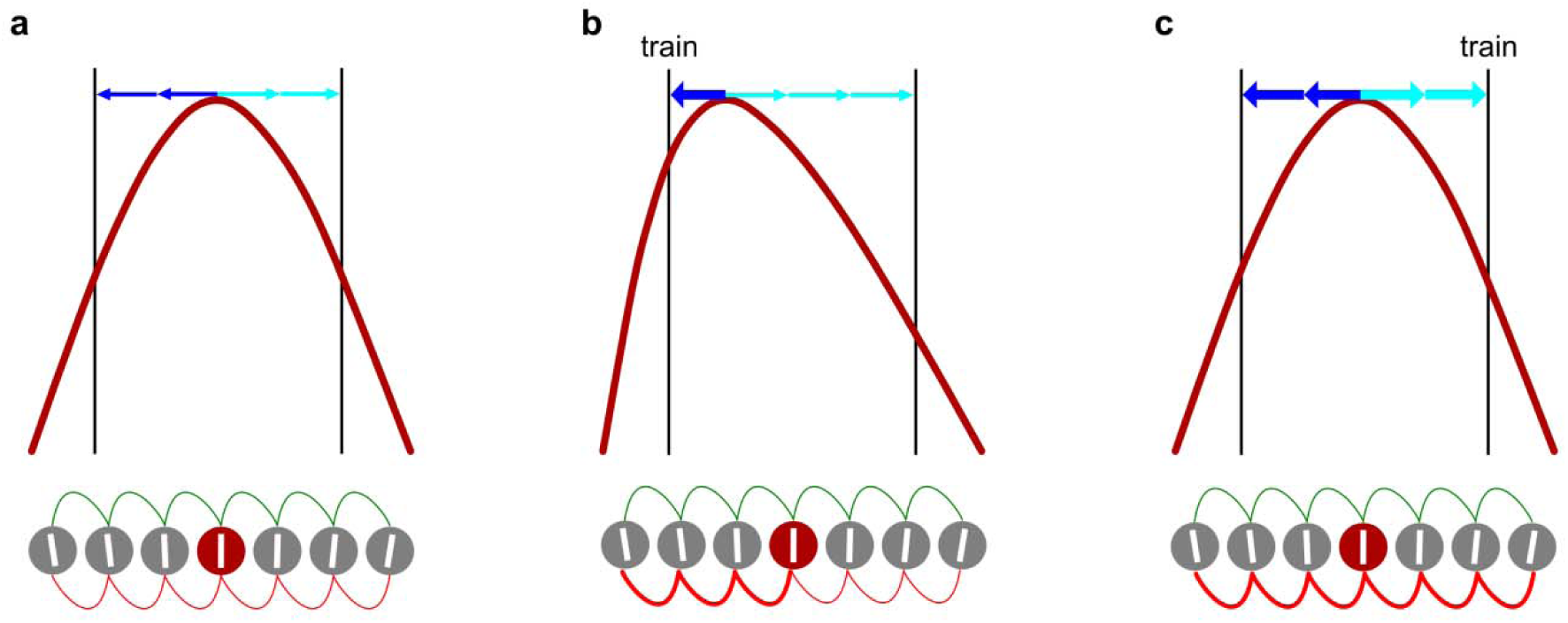
Schematic representation of synaptic and tuning curve changes due to training two similar orientations. **a**, A population of seven neurons (bottom row) with different preferred orientations (indicated by white bars). The red curve (top row) shows the tuning curve of the central neuron (red circle). The tuning curves of other neurons (gray circles) are omitted for clarity. Black vertical lines indicate two reference orientations at which training may occur. Tuning of each neuron results from broadly tuned feedforward input and excitatory (green) and inhibitory (red) recurrent connections with other neurons in the population. Recurrent network interactions effectively pull the tuning curve of the central neuron equally towards the left (blue horizontal arrows) and the right (turquoise horizontal arrows). **b**, Orientation discrimination training at the left reference strengthens inhibitory connections among neurons in the left portion of the network (red connections become thicker). This alteration of the recurrent interactions in the network effectively leads to increased pulling of the central neuron’s tuning curve towards the left reference (increased thickness of blue horizontal arrow). **c**, Orientation discrimination training at the right reference strengthens inhibitory connections among neurons in the right portion of the network. This alteration of the recurrent interactions in the network effectively leads to increased pulling of the central neuron’s tuning curve towards the right reference. This, in turn, rebalances the central neuron’s tuning curve. While two training events may thus lead to accumulation (rather than erasure) of network changes, they may cancel their respective effects on the shape of tuning curves.

Although considering learning and interference in terms of neuronal tuning correlates is instructive, it does not address how interference works at the level of memory traces embedded in a neural network as patterns of connectivity strengths. If there were interference, one possibility is that training in similar tasks requires incompatible network connectivity in the same population of connections, such that training at one reference orientation would erase the memory trace resulting from training at the other reference orientation. This may explain findings in a different experimental design reported by Ni and Maunsell (2010), and fits with ideas of ‘catastrophic interference’ in connectionist models of cortical learning due to different tasks posing different requirements on the same set of connections (McClelland, McNaughton, & O’Reilly, 1995). However, our network simulations suggest a radically different possibility, in which interference occurs without erasing any prior synaptic changes.

Our neural network was inspired by Teich and Qian (2003) who simulated orientation tuning in V1 by implementing thalamic feedforward input with broad orientation tuning which is sharpened by recurrent excitatory and inhibitory synapses among neighboring V1 neurons. We modeled training-induced tuning changes by strengthening inhibitory recurrent connections among orientation selective neurons in response to behavioral errors according to an error-triggered Hebbian learning rule. The choice to increase inhibitory connections, rather than reduce excitatory connections in line with an anti-Hebbian learning rule (Koch, Ponzo, Lorenzo, Caltagirone, & Veniero, 2013), is based on findings that learning changes the selectivity of inhibitory interneurons in V1 (Khan et al., 2018). In the model, illustrated in figure 1 from the perspective of a single example neuron with an orientation tuning curve shown in red, inhibitory and excitatory recurrent interactions are balanced prior to learning, yielding standard orientation tuning curves (figure 1a). Training at a reference orientation overlapping with one flank of the example tuning curve triggers locally restricted strengthening of inhibition among neurons best-tuned to the reference orientation. The enhanced inhibitory interactions among these neurons (figure 1b, bottom) will induce a collapse of the flank overlapping with the trained reference in the example tuning curve, combined with a tendency for the peak to be pulled towards the trained reference, and the opposite flank to become shallower. By analogy, subsequent training with a reference orientation overlapping with the opposite (flattened) flank will be associated with a matching enhancement of local inhibitory interactions (figure 1c), pushing the peak orientation back to its original location and restoring the symmetry of the example tuning curve. The restoration of the tuning curve symmetry in figure 1c is associated with a loss of expertise carried by the asymmetry of the tuning curve in figure 1b. Importantly, in this model the loss of expertise (interference) is not due to erasure of inhibitory connections that represent the memory trace. Rather, it is due to enhanced inhibition in other parts of the network.

This model makes two main predictions. Firstly, extensive training in an orientation discrimination task does not make it immune to interference by another task (figure 1b,c). Secondly, the number of trials required to create the inhibitory imbalance that enables high expertise equals the number of trials required to fully repair the inhibitory balance, which then would cause full interference (figure 1c). In other words, the model predicts that when two tasks are trained that require incompatible tuning curve shapes, discrimination performance on the two tasks will depend on the balance of trials devoted to each task. Our data support these model predictions; and hence also the idea that one skill can interfere with an earlier acquired skill, without erasing the connections related to the earlier established skill.

## 2 Model simulations

### 2.1 Model of orientation discrimination

V1 orientation tuning was simulated using a recurrent model first described by Teich and Qian (2003). Briefly, the model assumes that weakly orientation tuned feedforward information entering V1 from subcortical regions is sharpened by recurrent excitatory and inhibitory synapses among V1 neurons. In accordance with the descriptions given in Teich and Qian (2003) the model consists of *N* neurons with their preferred orientations *θ* covering 180 degrees. The firing rate *R*(*θ*, □*, t*) of each neuron with preferred orientation *θ* and presented with stimulus orientation □ at time *t* is given by

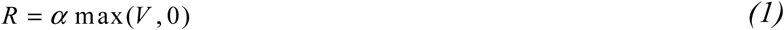

where *α* is a gain factor and *V* is the membrane potential which evolves according to

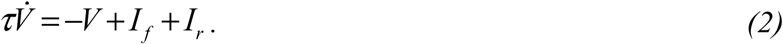

In equation 2, *τ* is the membrane time constant, *I_f_* the feed-forward input to the neuron, and *I_r_* the recurrent input a neuron receives from its neighbors. The feedforward input resulting from presenting stimulus orientation □ to a neuron with preferred orientation *θ* is given by

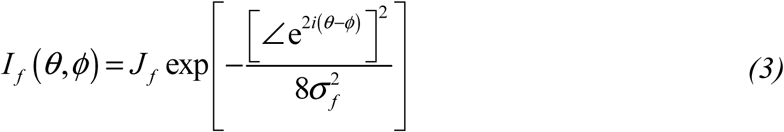

where *J_f_* and σ*_f_* respectively determine the strength and width of the input and the expression 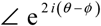 ensures that the phase difference wraps around the half circle. The recurrent input neuron *j* with preferred orientation *θ_j_* is the total activation it receives from neurons *i* with preferred orientations *θ_i_*

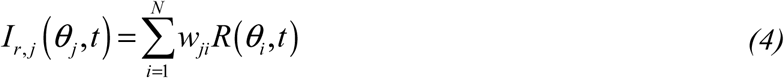

where *w_ji_* is the weight from neuron *i* with preferred orientation *θ_i_* to neuron *j* with preferred orientation *θ_j_*. The weight *w_ji_* is a composite of excitatory weight *w_exc,ji_* and inhibitory weight *w_inh,ji_* given by (*w_exc,ji_*-*w_inh,ji_*). Before learning, excitatory and inhibitory weights between two neurons *i* and *j* with respective preferred orientations *θ_i_* and *θ_j_* are initialized as

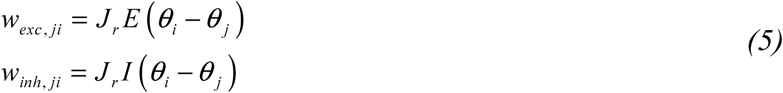

with *J_r_* controlling the connection strength and *E*(*θ*) and *I*(*θ*) being the excitatory and inhibitory connection probability distributions expressed by the periodic functions

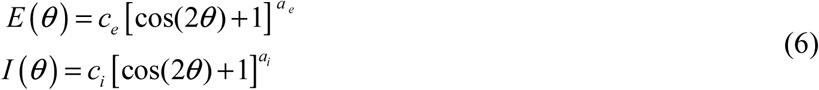

In equation 6, the exponents *a_e_* and *a_i_* control the sharpness of the excitatory and inhibitory distributions, respectively. The constants *c_e_* and *c_i_* are normalization factors ensuring that the sum of each distribution equals unity.

### 2.2 Behavioral performance and learning

In order to compare the performance of the model with that of our participants, the distribution of firing rates across neurons was translated into a decision of whether the currently presented orientation was rotated clockwise or counterclockwise with respect to the reference orientation. This was done using signal detection theory (Green & Swets, 1966). Specifically, the mean response of a neuron with preferred orientation *θ* to a stimulus with orientation □ is a function of the firing rate this stimulus evokes in the neuron *R*(□) and the duration of stimulation Δ*t*

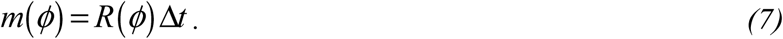

Furthermore, the variance of the firing rate under identical stimulus conditions was proportional to the mean response

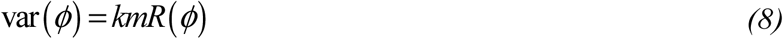

where *k* is a dimensionless constant between 1 and 4 (Peres & Hochstein, 1994; Shadlen & Newsome, 1994; Snowden, Treue, & Andersen, 1992; Softky & Koch, 1993). The activity *x*[*km*(□)] a neuron produces when presented with stimulus □ can be estimated by sampling from the normal distribution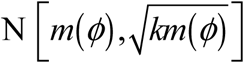. In order to make a decision whether a probe stimulus □_probe_ presented on a given trial was orientated clockwise or counterclockwise with respect to a reference stimulus □_ref_, each neuron calculated the difference of activity resulting from the two stimuli. Note that this implies that in the model, neural activity in response to the reference stimulus was explicitly calculated while participants were never explicitly presented with the reference stimulus. The difference [x(□ref) − x(□probe)] follows the normal distribution N 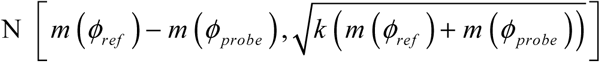. A neuron would make a correct decision either if it prefers □_ref_ over □_probe_ and if [*x*(□_ref_)−*x*(□_probe_)] *>* 0, or if it prefers □_probe_ over □_ref_ and if [*x*(□_ref_) *x*(□_probe_)] *<* 0. Taking these two cases together the probability that a neuron makes a correct decision (Green & Swets, 1966; Teich & Qian, 2003) is given by

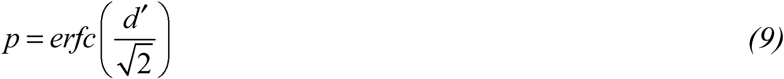

with

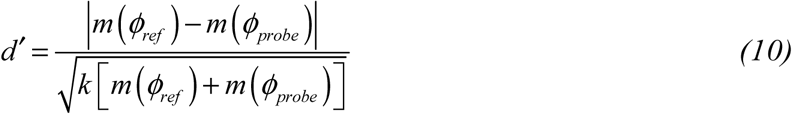

and *erfc*(·) being the complementary error function. A majority vote determined the performance of the population on a given trial, with a decision deemed correct if the percentage of all individual neurons making a correct decision exceeds a criterion *C*.

Learning, in our model, was a direct consequence of population performance within trials as it was implemented in the form of an error-triggered Hebbian learning rule affecting inhibitory connections (King, Zylberberg, & DeWeese, 2013). That is, whenever the population made an incorrect decision, reflective of similar response patterns generated by stimuli □_ref_ and □_probe_, inhibitory weights between active neurons were strengthened in order to decorrelate them and sensitize the population to stimuli around the reference orientation (Khan et al., 2018; King et al., 2013). Specifically, after each incorrect decision the weights were updated as follows

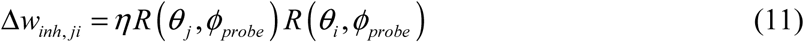

where *η* is the learning rate. After correct decisions the weights remained unaltered. Note that alternatively, we could have employed an anti-Hebbian learning rule, reducing excitatory weights after error trials (c.f. Koch et al., 2013). Mathematically these choices are identical in our model and lead to the same results (we confirmed this with simulations). Please also note that it is in principle possible that decorrelations occur additionally (or occur solely) after correct trials. Given an appropriately adjusted learning rate, such a scenario would not significantly affect results. However, correct trials imply that responses were already sufficiently dissimilar to allow for successful discrimination, rendering further decorrelation unnecessary. We therefore chose to let error trials drive the learning. Due to stochasticity in the decision-making process and hence in learning, all simulations were repeated 25 times to obtain good estimates of learning curves. The model is implemented in MATLAB (2015a, The MathWorks, Natick, MA) and freely available from https://github.com/ccnmaastricht/LTI.

Using the model we were able to reproduce learning curves reported for asymptotic perceptual learning in Been et al. (2011) by fitting three free model parameters to the data: a dimensionless constant *k* regulating the variance of neuron responses and hence affecting baseline performance, the positive learning rate *η* controlling the weight change in the Hebbian learning process, and a criterion *C* controlling the percentage of neurons that need to individually make a correct decision to deem the population response correct (for parameter values see table 1). Given the best fitting parameters, the root-mean-squared-error (RMSE) between model and empirical learning curves was 0.058.

**Table 1:** Parameters used in simulations.

| Symbol | Description | Value |
| --- | --- | --- |
| $N$ | Number of simulated neurons | 512 |
| $\tau$ | Membrane time constant | 15 ms |
| $\alpha$ | Gain of spike encoder | 10 spikes/s/mV |
| $J_r$ | Strength of recurrent connections | 1 mV/spikes/s |
| $J_f$ | Strength of feedforward connections | 0.5 mV/spikes/s |
| $\sigma_f$ | Gaussian width of feedforward orientation bias | 45° |
| $\eta$ | Learning rate | 1.4e<sup>-11</sup> |
| $k$ | Scaling of variance in the signal detection process | 1.05 |
| $C$ | Criterion for population decision | 53% |

## 3 Behavioral Experiments

### 3.1 Participants

Eight participants (mean age 23.98, *SD* 3.84, 6 female), naïve to the purpose of the study, were recruited. The sample size was based on a power analysis given effect sizes observed for interference training after a delay of 24 h in a previous study (Been et al., 2011; Cohen’s d ∼1.38), an *α*-level of 0.05, and a desired power of 0.95 for a one-sided repeated-measures t-test. All participants had normal or corrected-to normal visual acuity. Informed, written and verbalconsent wasobtained accordingtothe Helsinki Declaration, after full information about all procedures and the right to withdraw participation at any time. All procedures were approved by the local Ethical Committee of the Faculty of Psychology and Neuroscience (ECP). Participants received either monetary reward or credits to fulfill course requirements.

### 3.2 Stimuli, tasks and procedure

Stimuli used in the present study were Gabor patches (2.37 cycles/degree spatial frequency, 50% Michelson contrast, 56 cd/m^2^ average luminance, 3° diameter). They were presented on a gray screen (56 cd/m^2^) at 6° eccentricity from a centrally placed, white fixation dot along 45° or 135° polar angle lines, in one of three possible quadrants of the visual field. Participants were placed in a dimly lit room; their head was supported by a chin and head rest keeping eye-screen distance constant at 57 cm. The screen used was a 19 Samsung SyncMaster 940BF LCD monitor (Samsung, Seoul, South Korea; 60 Hz refresh rate, 1280*x*1024 resolution). The screen was covered by a gray mask with an oval aperture such that the screen borders were not visible to participants to prevent their utilization as reference for the orientation discrimination task. Fixation was monitored with a Viewpoint Eyetracker v.2.8.3 (Arrington Research, Inc., Scottsdale, Arizona, USA; 60 Hz sampling rate, 37 pixel/degree spatial resolution). Stimulus presentation and response recording was performed by Cortex v.5.9.6 (NIH freeware for psychophysical and neurophysiological experimentation).

Participants performed an orientation discrimination task with an unseen oblique orientation as reference stimulus. They indicated the direction of the orientation offset by pressing either the right or left arrow key for clockwise and counterclockwise rotations, respectively (figure 2). Hence, we employed a forced-choice identification design in which each stimulus required a ‘left’ vs ‘right’ decision. Each experimental trial started with a time window of maximally 750ms during which accurate fixation was to be initiated (i.e., deviation *<*1.5° from fixation dot) and subsequently maintained for another 250ms to trigger stimulus presentation as well as throughout presentation of the stimulus (500ms) in one of the four quadrants of the visual field. Stimulus presentation was then followed by a 1000ms response window. Responses were given with the right middle and index fingers indicating clockwise or counterclockwise deviations from the reference orientation, respectively. Participants received feedback after each trial in the form of color changes (green = correct; red = incorrect) of the fixation dot lasting 200ms. After feedback, the fixation dot disappeared signifying the end of the trial, followed by a fixed 500ms inter-trial interval. When a participant’s gaze fell outside the fixation window during the fixation period, trials were aborted. Aborted trials were repeated at a randomly chosen time during the experiment.

**Figure 2.**
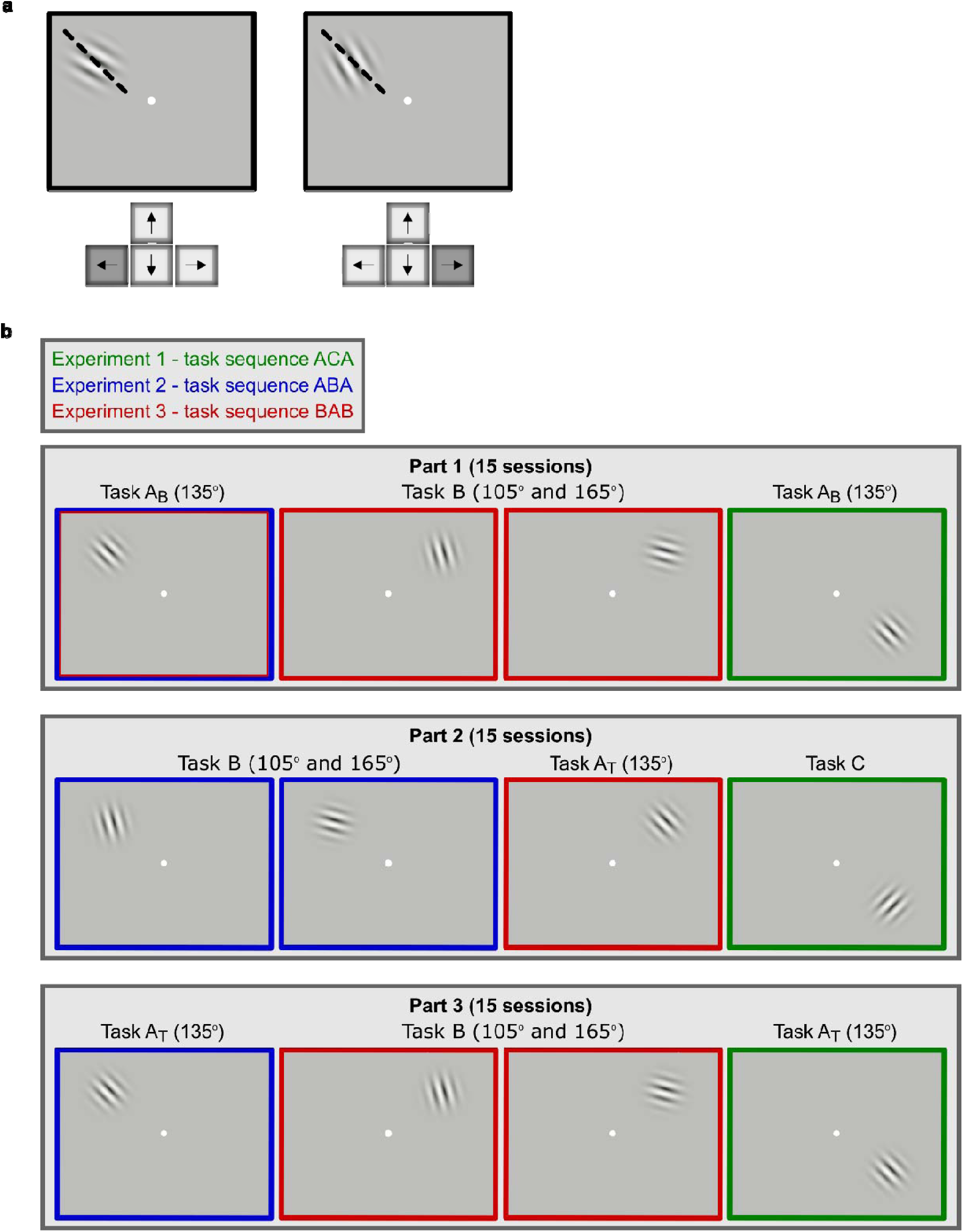
**(previous page): a**, Orientation discrimination task and stimuli. Thresholds were determined using an identification design, with left key responses for counterclockwise orientation offsets from a 135° reference (dashed line, not shown to participants), and right key responses for clockwise orientation offsets. Stimuli are enlarged for illustration. **b**, Design of the study. The study was split in three periods (Parts 1-3) and each part consisted of four conditions. Parts 1, 2, and 3 lasted 15 sessions each and were done consecutively (order of conditions within each part was counterbalanced over participants). In total, the study took 45 sessions. The conditions are named according to executed task, with Task A referring to orientation discrimination at reference orientation 135°, Task B referring to the combined training at 105° and 165°, and Task C referring to orientation discrimination at reference orientation 45°. A learning curve in a Task can serve as a Baseline against which the effect of interference is tested (indicated by subscript B in Task A_B_ and Task B_B_). A learning curve that has undergone the effect of interference represents a Test of interference (indicated by subscript T in Task A_T_ and Task B_T_) The frame color indicates conditions of stimuli presented at the same location, thus belonging to the same experiment, with green, blue, and red frames corresponding to Experiments 1, 2, and 3, respectively. The second frame in the upper row **e** is marked in both blue and red, to indicate that this condition provided a baseline measurement for Task A for both Experiment 2 (blue frames) and 3 (red frames).

In the present study, we conducted three experiments, each in a different quadrant of the visual field. Experiment 1 was a control experiment in which we tested whether 15-session training at a 135° reference orientation (Task A_B_) would be interfered with by subsequent 15-session training at an orthogonal reference orientation (45°; Task C), as tested in subsequent training at the 135° reference orientation (Task A_T_). Based on previous work (Been et al., 2011) and model simulations we did not expect interference. Experiment 2 was designed to test whether a 15-session baseline training at a 135° reference orientation (Task A_B_) would be interfered with by subsequent 15-session training at similar reference orientations (105° & 165°; Task B). This was tested in an additional 15-session period at the 135° referenceorientation (Task A_T_). Experiment 3 was designed to test whether a 15-session training at reference orientations 105° and 165° (Task B) would interfere with subsequent training at a 135° reference orientation (Task A_T_). In addition, Experiment 3 permitted testing whether Task A would interfere with subsequent learning in Task B_T_as compared to Task B_B_. The sequences of training at different reference orientations and the orientation differences among reference orientations in the different experiments were based on (Been et al., 2011).

Figure 2b shows how these three experiments were combined into a single study design. The study consisted of three parts, each lasting for 15 sessions. During each session, participants completed all four conditions belonging to a Part, with each condition representing training at a specific reference orientation andin a specific visual field quadrant location. For each condition (i.e. each frame in figure 2b), participants completed four staircases, such that they performed 16 staircase measurements per session. Participants spent 15 sessions performing Part 1, then 15 sessions performing Part 2, and then 15 sessions performing Part 3. Figure 2b also makes clear that the assignment of conditions to numbered experiments is conceptual, and unrelated to the time period in which the data were collected. After performing the 45 sessions, all three planned experiments were completed, and analysis was based on three separate selections of conditions representing the three experiments (color-coded in figure 2b with green = Experiment 1; blue = Experiment 2; red = Experiment 3).

In the execution of the study, there were two versions that determined in which visual field location each of the three experiments were performed. In version A (illustrated in figure 2b), Experiment 1 was done in the right lower quadrant, Experiment 2 in the left upper quadrant, and Experiment 3 in the right upper quadrant. Version B was mirror-reversed relative to version A along the vertical meridian. Condition order within a Part was counterbalanced across days/participants. In each session, irrespective of the part of the study, participants performed the task on oblique stimuli (condition A using 135°, and C using 45°) before they performed the task on flanking orientations (condition B, using 105° and 165°). The order of the two conditions using oblique orientations was counterbalanced within participants, as was the order of conditions using flanking orientations. Within participants, testing schedules were kept constant (i.e., at least three sessions per week, all at the same time of day) for the duration of the study. Importantly, although each Part comprised 15 daily sessions, the calendar length for the full training in each Part comprised 4 weeks (mean 28 days, SD 7.34 days), as there was no testing on weekends, and as participants typically could be tested only 3 to 4 times per week. The whole study in each participant therefore lasted 3 months (mean 90 days, SD 19.06 days on average; see table 2).

**Table 2:** Duration of experiment for individual participants.

| Prt | P1 | $\Delta(1-2)$ | P2 | $\Delta(2-3)$ | P3 | total |
| --- | --- | --- | --- | --- | --- | --- |
| 1 | 25 | 2 | 25 | 0 | 34 | 86 |
| 2 | 29 | 5 | 29 | 0 | 36 | 99 |
| 3 | 33 | 0 | 18 | 26* | 52 | 129 |
| 4 | 24 | 0 | 21 | 0 | 26 | 71 |
| 5 | 29 | 0 | 38 | 1 | 25 | 93 |
| 6 | 21 | 0 | 19 | 2 | 25 | 67 |
| 7 | 35 | 0 | 27 | 0 | 23 | 85 |
| 8 | 32 | 0 | 26 | 0 | 31 | 89 |

Duration of the different training conditions and the total experiment for each participant (Prt). The number of days for each training part (1, 2, 3) in days are shown in columns 'P1', 'P2', and 'P3', respectively. The number of days without any training in between training parts is indicated in columns ' $\Delta(1-2)$ ' and ' $\Delta(2-3)$ '. \* This break was longer due to closure of the university building during Christmas holidays.

### 3.3 Threshold measurements

We used a Wetherill staircase tracking 84 % correct performance to acquire just-noticeable differences (JND; Wetherill & Levitt, 1965). The staircase ended when either 14 reversal points were acquired or 120 trials were completed. In total, each participant completed 16 staircases in each session, and a total of 720 staircases (or ∼72000 trials) divided over 45 sessions. From each staircase, the last ten reversal points were selected to contribute to threshold estimation. Since four staircases were done per quadrant in each session, we determined the threshold per quadrant/session as the geometric mean of 40 reversal points.

Note that the threshold corresponds to the difference in the stimulus orientations that participants could discriminate at 84% correct, and not the absolute deviation in angle between the stimuli and the theoretical reference orientation as done in a number of other studies (Schoups et al., 1995, 2001). Our thresholds would have to be divided by two to compare them directly with these studies. On the other hand, our thresholds are directly comparable to those reported in other studies (Raiguel et al., 2006; Yang & Maunsell, 2004). In the first session, the start level of each staircase was set to an orientation difference of approximately 15°, which corresponds to the 84% orientation threshold reached on average by naïve participants in our task (see Been et al., 2011). In participants for whom this difference is too small, a few error responses will immediately lead to much larger differences that can serve as examples. For all subsequent sessions, the average threshold of the previous session was taken as starting level. The orientation difference was adapted to performance in a proportional manner, by either dividing or multiplying the current orientation difference by a factor of 1.2, depending on performance criteria designed for the staircase to converge on an 84 % correct level. All thresholds were ln-transformed and are publicly available (https://doi.org/10.5061/dryad.6djh9w0wn).

### 3.4 Statistical analysis

We used repeated measures t-tests to assess whether the mean difference between ln-transformed JNDs observed for asymptotic performance before and after interference training were significantly different from zero. In order to assess whether mean difference between JNDs observed for two conditions were equivalent, we used two one-sided tests of equivalence for paired samples (TOST-P; Seaman & Serlin, 1998).

While the TOST-P, and equivalence testing in general, cannot establish exact correspondence between two conditions, it can establish that differences fall within a pre-determined interval reflecting negligible effect sizes. Specifically, the TOST-P specifies a lower and an upper equivalence bound based on the smallest effect size (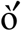) of interest and performs two t-tests on the Null hypotheses 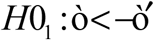 and 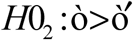. If both Null hypotheses can be rejected, the observed effect lies within the interval 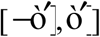 and is considered sufficiently small to be irrelevant. In line with a procedure outlined in (Mara & Cribbie, 2012), taking sample size and variance of a specific data set into account, we constructed an equivalence interval by first specifying a negligible difference and constructing a 95% confidence interval around this difference. We regard a difference of 0.31 log units (∼1.36°) sufficiently small to be negligible. We obtained this value by examining the extremes of expected differences in identical conditions. This was done by performing a blocked bootstrapping procedure. From N participants, we repeatedly (100,000 times) sampled (with replacement) two learning curves (each comprising 15 sessions) obtained separately in each participant in equivalent conditions A and B. Prior to drawing each sample, we randomly assigning the labels A or B to the two curves of each participant. We then computed the performance difference between samples A and B either for early (first three sessions) or asymptotic learning (last three sessions) for each sample. We considered the 99^th^ percentile of the distribution of absolute performance differences to be a good indicator of negligible differences given that conditions are indeed equivalent. We performed this procedure using two sets of data. First, from the data set gathered for this study, we used learning curves for baseline training at the 135° reference in Experiments 1 and 2. Second, we used data from a previous study (Been et al., 2011). Specifically, we used learning curves for training at a 135° reference in one of the two upper visual field quadrants and the average of learning curves for training at 105° and 165° references in the other upper visual field quadrant. On average (28 participants, one excluded because of lacking data), these 15-session learning curves were near-identical (for a discussion of that result, see Been et al., 2011). From our own dataset we obtained estimates of negligible difference for early learning (0.29 log units) and late learning (0.33). From Been et al.’s (2011) data, we obtained similar estimates for early learning (0.27) and late learning (0.35), respectively. Given the similarity of these values we used the average of these four values as a single estimate of equivalence (0.31). To construct the confidence interval, we estimated the standard error of the differences between the empirical baseline learning curves in our Experiments 1 and 2 at the beginning and end of the learning curves (which was 0.29 on average). This led to a final equivalence interval of [−0.60,0.60] in log units, corresponding to [−1.83°,1.83°]). We thus consider two JNDs that differ by 1.83° or less to be equivalent in the context of our experimental paradigm. Goodness of fit between simulated and empirically observed asymptotic learning curves was measured by their root-mean-squared-error (RMSE).

## 4 Results

### 4.1 V1 tuning curve changes underlying perceptual learning and behavioral interference

Because V1 neurons show orientation tuning (Hubel & Wiesel, 1959; Schoups et al., 2001) and are retinotopically organized (Daniel & Whitteridge, 1961; Gattass, Sousa, & Rosa, 1987), similarly oriented stimuli presented at the same visual field location will stimulate overlapping neuronal populations, thus creating the preconditions for potential behavioral interference. To enhance the relevance of V1 properties for hypothesis testing, we trained participants to reach asymptotic levels of performance, during which the contribution of V1 is increased (Hochstein & Ahissar, 2002; A Karni & Bertini, 1997; Schoups et al., 2001). Accordingly, the V1-inspired computational orientation discrimination model we used for making quantified predictions of learning (and interference) was restricted to asymptotic learning.

In our V1 model, tuning curves resulted from broadly tuned feedforward input sharpened by recurrent excitatory and inhibitory connections (Teich & Qian, 2003), in line with known mechanisms of orientation tuning (Ferster, Chung, & Wheat, 1996; Shapley, Hawken, & Ringach, 2003; Sillito, 1975). Orientation tuning enabled the model to make decisions regarding the rotation of an oriented stimulus relative to a reference orientation based on differential responses to the two orientations. To simulate skill learning, we allowed the model to learn from feedback regarding the correctness of its decisions, by adjusting inhibitory recurrent connections according to an error-triggered Hebbian learning rule (Khan et al., 2018; King et al., 2013; see section 2.2 for details). The resulting shift in the excitatory-inhibitory ratio of recurrent connectivity led to sharpening and flattening in the flanks of model neuron tuning curves in accordance with empirical studies showing lasting tuning changes in low-to-mid-level visual areas after long-term asymptotic learning in difficult discrimination tasks (Raiguel et al., 2006; Schoups et al., 2001; Yang & Maunsell, 2004). These tuning changes are driven by the fact that the flanks of tuning curves are most informative in signaling small changes in stimulus orientation (Raiguel et al., 2006; Schoups et al., 2001). Free model parameters were calibrated based on human asymptotic learning curves for orientation discrimination from a separate study (Been et al., 2011; see section 2 for details). Figure 3a illustrates the naïve state of the network model by example tuning curves with preferred orientations of 90°, 105°, 120°, 135°, 150°, 165°, and 180°. These tuning curves were equally spaced, identical and symmetric. Figure 3b shows the expected network state after extensive training at a 135° reference orientation (Task A). Tuning curve flanks that overlap with the trained reference orientation show training-induced sharpening (see blue tuning curves in figure 3b).

**Figure 3.**
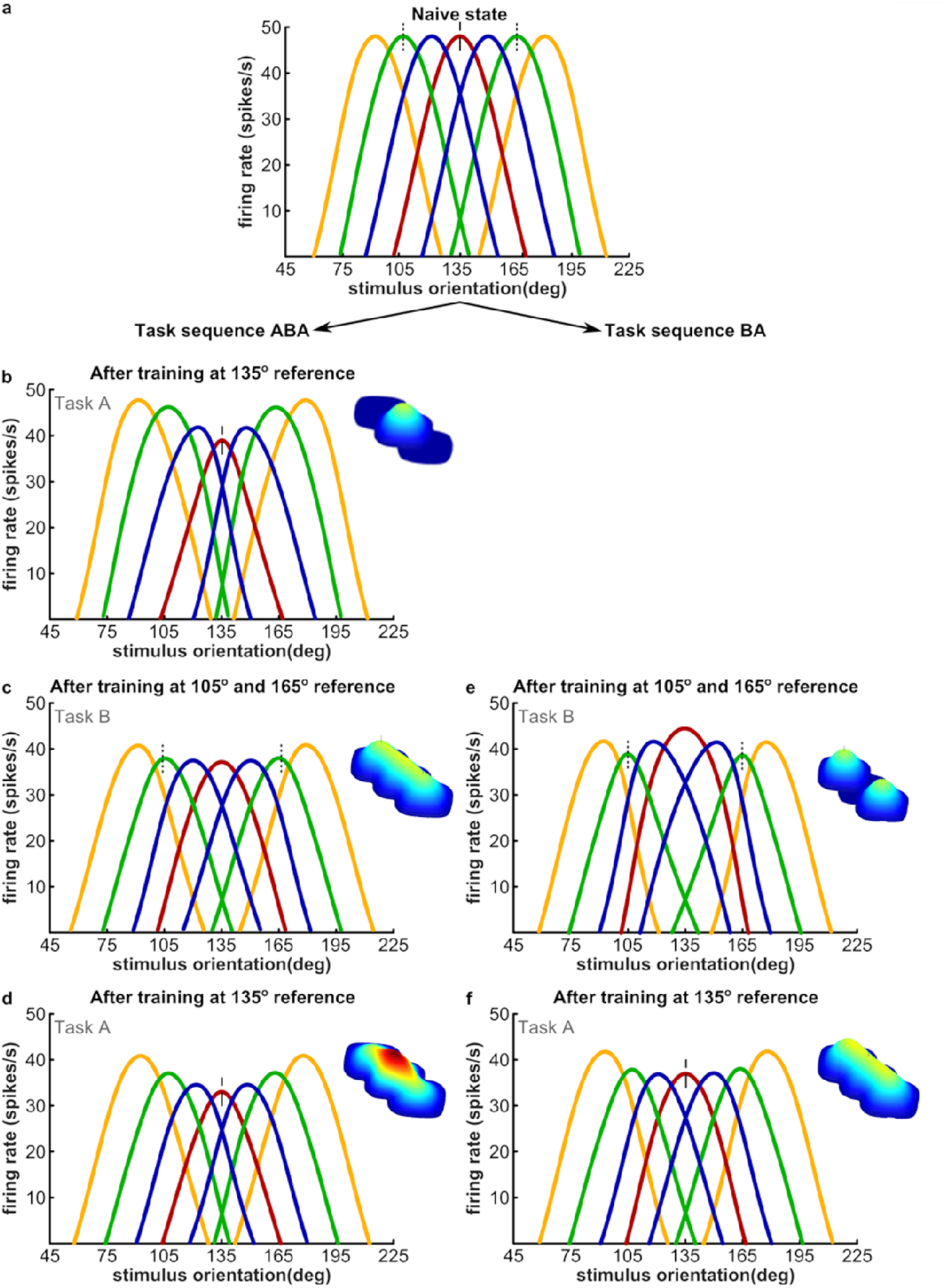
**(previous page):** Modeling of orientation tuning changes at the end of asymptotic learning in a V1-like cortical area resulting from different learning sequences. Training-induced tuning changes are exaggerated for illustrative purposes. Panels **b-d** represent consecutive periods of prolonged training at 135° (Task A), at 105° and 165° (Task B), and finally again at 135° (Task A). Likewise, panels **e-f** represent consecutive periods of extended learning, first at 105° and 165° (Task B), and then at 135° (Task A). Thin vertical lines at top of tuning curves indicate the position of the reference orientations (dashed: 135° reference; dotted: 105° and 165° references). **Insets** in panels **b-f** graphically depict the size of networks contributing to orientation discrimination of the three reference orientations. Training-induced enhancement of inhibition is shown by a color-coded height of distribution on top of the network (saturated red = largest increase in inhibition). **a**, Orientation tuning curves prior to any learning. **b**, Network state and tuning curves after 15 training sessions at the 135° reference orientation. Neurons with preferred orientations ± 15° away from the trained reference show an increased slope of the flank overlapping with the trained reference (blue curves), paired with the tops shifting somewhat towards the trained reference. Increasing the slope overlapping with the trained reference is useful in generating larger differential responses for small orientation changes around the reference. The inset distribution displays an increase in inhibitory weights with respect to the naïve network state. All changes are clustered around 135°. **c**, Network state and tuning curves after training at the 105° and 165° reference orientations. This pulls the blue curves back into place, generating a distribution relatively similar to the naive state. Thus, Task B almost ‘resets’ all neuronal tuning curves crucial for learning Task A, and effectively erases most of the skill at the 135° reference orientation built up during initial learning. Changes in recurrent connectivity due to increased inhibition now extend to 105° and 165° (light blue-yellow regions in inset). Thus, synaptic changes around 135° remain and merge with new changes. Note that the state reached after training Task A (panel **b**) does not prevent the learning in Task B at reference orientations 105° and 165° (panel **c**). Training in Task A flattens the tuning curve slopes contributing to learning at the 105° reference in about half of the neurons (blue curves), but not the other half (orange curves). The same is true for the 165° reference. Hence, Task A leaves intact about half of the neuronal tuning curves relevant for learning Task B. Therefore, no or only limited interference of Task A upon Task B is expected. **d**, Restoration of skill at the 135° reference by new training, following interference by 105° and 165°. Lateral inhibition around 135° exhibits additional increases (red color in inset). **e**, Network state after an initial period of extensive training at the 105° and 165° references. **f**, Network state after extensive training at the 135° reference when preceded by extensive training at the 105° and 165° references. Failure to reach the fully trained state as in panel **b** corresponds to predicted interference.

Extending the reasoning based on figure 1, expertise accumulated at 135° (Task A) should not be interfered with by following training at a 45° reference orientation (Task C). The limited width of tuning curves ensures that training in the same location at these two reference orientations would be based on non-overlapping neural populations. To test the absence of interference when subsequent training periods use orthogonal reference orientations, the model was subjected to a training sequence consisting of 15 sessions Task A, followed by 15 sessions Task C, and 15 sessions Task A (ACA, Experiment 1). Model simulations confirmed the absence of interference (not illustrated in figure 3).

Based on figure 1, an effective way to interfere with the expertise accumulated at reference orientation 135° (Task A) would be to train at both the 105° and 165° reference orientations (together referred to as Task B). Hence, we exposed our model to 15 sessions of training in Task A, followed by 15 sessions Task B, and another 15 sessions Task A (ABA, Experiment 2, figure 3b-d). Note that Task B will always recruit the full neural population storing skill for Task A, but that Task A will recruit only about half of the neural population storing the skill for Task B (see legend figure 3 for explanations). Hence, while we expected Task B to interfere with Task A, Task A was expected to be relatively ineffective in interfering with Task B (Vogels, 1990).

A detailed description of model simulations for sequence ABA is shown in Figures 3b-d. After asymptotic training at reference orientation 135° (Task A), neurons originally tuned to orientations deviating by ∼+ 15° or ∼−15° from the trained reference (blue tuning curves figure 3), steepened their flanks facing the trained reference orientation and moved their tops towards the reference. This, in turn, resulted in larger differential responses to similar orientations thereby enabling better discrimination performance. The training-induced asymmetry of the tuning curves reflects an imbalance of lateral inhibition (see also figure 1). For example, the changes for the blue tuning curve tuned to ∼150° (figure 3b) stemmed from an imbalance in lateral inhibition in the network providing input to the 150°-tuned neuron. Due to a training-induced increase in lateral inhibition among neurons with preferred orientations in the 135°-150° range, inputs to the 150°-tuned neuron from precisely those neurons that were less informative for discriminations around the 135° reference (red tuning curve in fig 3b) were diminished. At the same time, the influence from neurons with peak response to orientations >150°, whose role in the task was limited, remained largely unaltered, thus creating the inhibition imbalance. Analogous reasoning applies to neurons with tuning around 120°. A simulation of subsequent asymptotic learning at 105° and 165° reference orientations (Task B, figure 3c) reversed the steepening effects that took place in preceding learning and moved the tuning curve maxima back to their original places, restoring the population to a quasi-naive distribution of tuning curves. Hence, when baseline training of the model at 135° (Task A) was followed by training at 105° and 165° (Task B), original learning in Task A was erased. Importantly, synaptic changes in response to initial training on Task A were not undone by subsequent training on Task B. Rather, training-induced lateral inhibition increases around the 135° reference after Task A were matched by an equal increase in lateral inhibition near references 105° and 165° after Task B (figure 3c, inset shows the distribution of inhibitory weights). The acquisition of two memory traces thus rendered both of them inconsequential due to the re-balancing of lateral inhibition. Therefore, another period of asymptotic learning at 135° was required to approach restored performance levels at this reference orientation (figure 3d). Thus, simulations confirmed the effectiveness of Task B to interfere with Task A. Simulations also confirmed that Task A was rather ineffective in interfering with Task B. The state of the population after training on Task A (figure 3b) did not prevent effective learning of Task B. This is due to the fact that training in Task A left about half of the neuronal tuning curves critical for the learning of Task B unaffected (orange curves in figure 3b).

Finally, we describe model simulations for sequence BA. Training in Task B (at 105° and 165°) created a state (figure 3e) that was highly unfavorable for following training in Task A (at reference orientation 135°). In this case, training at 135° in Task A matched in duration to Task B may at best undo the detrimental effects of preceding training in Task B by reverting the tuning curve distribution back to a quasi-naïve state (figure 3f). Behavioral interference according to our model is thus the result of competing demands placed on tuning curve changes. The incompatibility of tuning curves in the neural network can be appreciated by comparing figures 3b and 3e. Note additionally that enhanced inhibition in subparts of the network after training Task B, is rebalanced by increased inhibition in an additional part of the network (insets in figure 3e and f).

Note that in our model, training at a reference orientation leads to reduced responses in neurons with preferred orientations at or near the reference orientation (e.g., compare figures 3a-d). This is in line with a possible link between adaptation and learning as suggested in empirical (Dragoi, Sharma, & Sur, 2000) and modeling studies (Teich & Qian, 2003). In our model, the response decreases reflect enhanced lateral inhibition after Hebbian learning (see insets in figure 3). Note also that when a new period of training follows a previous period, (e.g., figure 3c and 3f) the strengthening of inhibitory connections in a network subregion due to the new training could marginally overlap and add to the pre-existing strengthened inhibitory connection weights set by the prior training. Furthermore, re-learning a task after interference (figure 3d) involves re-creating an inhibition imbalance by adding inhibition to the network subregion on top of pre-existing inhibition from prior training in the same task.

### 4.2 Psychophysical demonstrations of interference between skills trained in 4-week long, successive training periods

We trained eight human participants on an orientation discrimination task using a Gabor stimulus of 3° diameter presented at 6° of eccentricity. They were exposed to the same experiments as our model. Making use of position specificity of orientation discrimination at least at the level of visual field quadrants (Been et al., 2011; De Weerd, Peralta, Desimone, & Ungerleider, 1999; Schoups et al., 2001), three experiments (task sequences: ABA, BAB, ACA) were distributed over different quadrants, allowing us to test interference between the various tasks within each participant. Per experiment, tasks Aand B (or C) were trained in three consecutive periods each lasting 15 sessions (spread in each participant over 3 months in total). We tested whether skill buildup in Task A (135° reference) during baseline training (referred to as Task A_B_) suffered from training in Tasks B (105°/165°) and C (45°) by testing (and training) again in Task A (referred to as Task A_T_). We also tested whether Task B suffered from prior training in Task A. Within each session, participants performed four 84 % correct threshold measurements per combination of task and quadrant (see section 3 for details).

In the statistical evaluation of the psychophysical data, we considered effects of interference on late learning (last three sessions of a learning curve) as well as on early learning (first three sessions). While our model can be used to predict effects of late learning, we have no model at present that predicts effects of interference on early learning. A tentative interpretation of any interfering effects on early learning is deferred to the Discussion.

In Experiment 1, using task sequence ACA (figure 4a-c), we did not expect interference of Task C on Task A. Our results showed a trend for the asymptote in Task A_T_ to converge on smaller thresholds than the asymptote in Task A_B_. A planned one-sided t-test comparing later learning in Tasks A_B_ and A_T_ only marginally failed to show this difference (*t*_(7)_ = - 1.65, *p* = 0.07). In addition, the mean performance levels at the start of Task A_T_ (sessions 31-33) and the end of Task A_B_ (sessions 13-15) were significantly equivalent (*t*_1(7)_ = - 7.251, *t*_2(7)_ = 4.670, *p* = 0.01). Together, these analyses confirmed that skill at 135° in Task A was not interfered with by training at an orthogonal reference orientation in Task C. Instead, there was a trend towards additional learning in Task A_T_.

**Figure 4:**
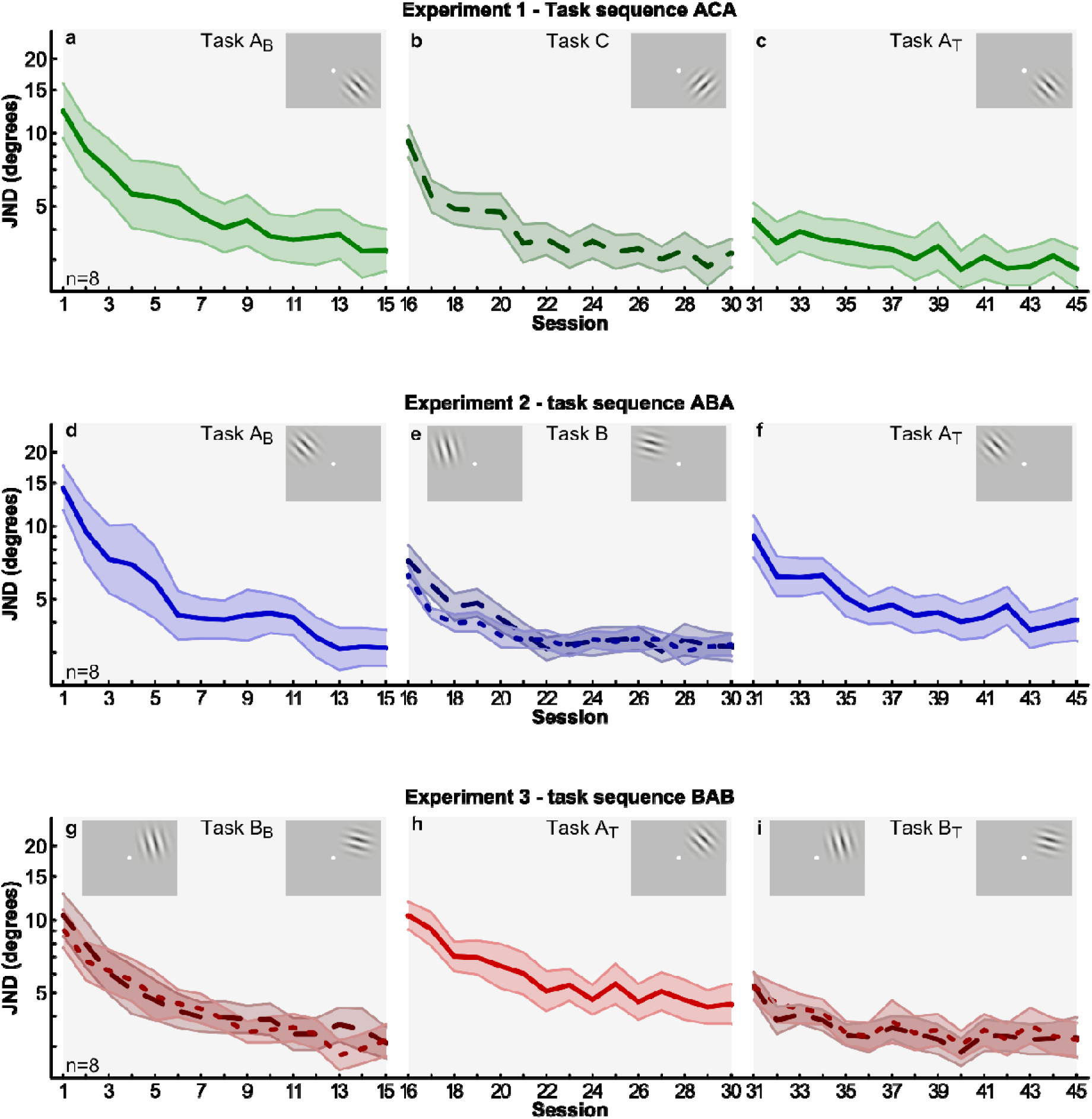
Learning curves in three different sequences of learning periods averaged over eight participants. A period consisted of 15 sessions of orientation discrimination training spread over a month in Task A (reference 135°), Task B (references 105° and 165°) or Task C (reference 45°). When Task A was trained first, it is referred to as a ‘Baseline’ (Task A_B_), and when it was trained last, it is referred to as a ‘Test’ (Task A_T_). An analogous nomenclature is used for Task B. We refer to the rows as Experiments 1 to 3 (top to bottom). A total of nine conditions were performed by each participant, in a semi-random order illustrated in figure 2 in section 3.2. Colors of curves correspond to conditions in figure 2b. **a-c**, Thresholds as a function of training session in Experiment 1 (Task sequence ACA). **d-f**, Learning curves in Experiment 2 (Task sequence ABA). **g-**i, Learning curves in Experiment 3 (Task sequence BAB). In Task B, the coarser dash corresponds to the 105° reference orientation. Insets schematically illustrate the different task conditions (stimuli enlarged for illustration). Columns indicate the temporal order in which conditions were trained (with the conditions in the left column trained in the first month, those in the middle column in the second month, and those in the right column in the third month; see section 3.2, Methods).

In Experiment 2, participants were trained in task sequence ABA (figure 4d-f). Qualitatively, our model simulations for this sequence (figure 3b-d) predicted that in spite of extensive baseline training in Task A_B_ at the 135° reference orientation, the intervening training in Task B at 105° and 165° would interfere with performance in Task A_T_ in the test period (figure 4). Planned t-tests revealed that mean performance at the end of Task A_T_ (sessions 43-45) and the end of Task A_B_ (sessions 13-15) were significantly different (*t*_(7)_ = 2.144, *p=* 0.0346). In addition, performance thresholds at the end of Task A_T_ in Experiment 2 were substantially elevated beyond those observed for Task A_T_ in Experiment 1 (compare figs 4c and 4f; *t*_(7)_ = 3.823, *p* = 0.0065). Hence, Task B interfered with late learning of Task A.

In addition, we found in Experiment 2 that average thresholds at the beginning of learning in Task A_T_ (sessions 31-33) were significantly elevated above the end of asymptotic learning in Task A_B_(sessions 13-15; *t*_(7)_ = −5.587, *p* < .001). We then tested whether performance at the start of Task A_T_ (sessions 31-33) was the same as that at the start of Task A_B_ (session 1-3). However, two one-sided tests of statistical equivalence for paired samples with an equivalence interval of ([−1.83°,1.83°]) failed to show that these performance levels were significantly equivalent (*t_1(7)_* = −1.155, *t_2(7)_* = 4.186, *p* = 0.143). Please note that failure to reach significance in the equivalence test does not permit the conclusion that performance levels are different. Indeed a post-hoc two-sided t-test failed to establish any significant difference between these performance levels (*t*_(7)_ = 1.62, *p =* 0.149). Hence, thresholds at the beginning of the learning curves for Task A_B_ and Task A_T_ were neither significantly equivalent nor significantly different. In summary, Task B interfered with the beginning of learning in Task A, in addition to interfering with late learning in that task.

In Experiment 3, participants followed task sequence BAB (figures 4g-i). For the BA part of the sequence, we expected that Task B (figure 4g) would proactively interfere with Task A (figure 4h). In accordance with an interference effect (see also figure 3e-f), a one-sided t-test showed that thresholds at the end of asymptotic learning at 135° in Task A_T_ (sessions 28-30, figure 4h) were significantly elevated compared to those at the end of asymptotic learning in the baseline condition (sessions 13-15; *t*_(7)_ = 2.9238 *p* = .011; figure 4d). These data support substantial interference of Task B upon asymptotic learning in Task A.

In a secondary analysis, we also investigated in Experiment 3 whether performance in Task A (figure 4h) interfered with previously built up skill in Task B_B_, and thus would interfere with subsequent training in task B_T._ We did not expect such interference. To test this, we compared thresholds at the end of learning curves in Task B_T_ (figure 4i) and Task B_B_ (figure 4g), and found them to be statistically equivalent (for 105° *t*_1(7)_ = −3.838, *t*_2(7)_ = 4.763, *p =* 0.003; for 165° *t*_1(7)_ = −6.372, *t*_2(7)_ = 4.170, *p =* 0.002). We further tested whether performance at the beginning of Task B_T_ experienced interference from training of Task A. As expected, a one-sided t-test revealed that thresholds at the beginning of Task B_T_ (sessions 31-33) were significantly lower than those at the beginning of Task B_B_(sessions 1-3), despite intervening training on task A (*t*_(7)_ = - 3.523, *p =* 0.005 & *t*_(7)_ = −2.340, *p =* 0.026 for reference orientations 105° and 165°, respectively). This contrasts to the analogous comparison for Experiment 2, where the beginning thresholds for Tasks A_B_ and A_T_ were difficult to distinguish. Finally, we tested whether mean performance levels at the start of Task B_T_ (sessions 31-33) were significantly equivalent to those at end of Task B_B_ (sessions 13-15). For the 105° this was indeed the case (*t*_1(7)_ = −6.167, *t*_2(7)_ = 2.709, *p =* 0.015). For 165°, surprisingly, the corresponding performance levels narrowly failed to reach significant equivalence (*t*_1(7)_ = - 9.913, *t*_2(7)_ = 1.415, *p =* 0.1). A post-hoc t-test revealed that these performance levels were instead significantly different (*t*_(7)_ = −4.542, *p =* 0.003). The intervening training in Task A thus had some, albeit small, effects upon beginning thresholds in Task B_T_. In sum, as expected, this secondary analysis in Experiment 3 showed that Task A did not interfere with asymptotic learning in Task B, and at best interfered only weakly with the initial learning in Task B.

When putting together the evidence from Experiments 2 and 3, we found that Task B (as expected) showed a much stronger capacity to interfere with Task A than vice versa. Specifically, with respect to late learning, Task B strongly interfered with Task A, but Task A did not interfere with Task B. These results fit with a contribution of V1 (and other low-level visual areas) to skill memory formation during late learning. The results on late learning confirm that unless neural overlap between tasks is limited by choosing stimuli encoded by distinct neural populations, behavioral interference will occur. With respect to early learning, Task B strongly interfered with early learning in Task A, whereas Task A produced no or only weak and inconsistent interference on Task B. The interference effects involving early learning may thus involve mechanisms that go beyond early visual cortex (see Discussion).

In agreement with other studies (De Weerd et al., 1999; Raiguel et al., 2006; Schoups et al., 1995, 2001), we found that orientation discrimination learning is specific for reference orientation and position (e.g., compare learning curves during Part 1 of training, left hand column in figure 4). This predicts that the different interference effects also remain specific for the quadrants in which they are elicited (Been et al., 2011). The third Part of training (right hand column figure 4, sessions 31-45) lent itself to a statistical test of the location specificity of interference. In the third Part, we compared performance in Task A_T_ in one location exposed to training sequence ABA (figure 4f) to another location exposed to training sequence (ACA) (figure 4c). If interference were position specific, threshold measurements in Task A_T_ are expected to be elevated when preceded by Task B (figure 4e), but not when preceded by Task C (figure 4b). These predictions were confirmed by a one-sided t-test both for early learning (sessions 31-33, *t_(7)_* = −9.060, *p* < 0.001), and for late learning (sessions 43-45, *t_(7)_*= −3.823, *p* = 0.003), thus indicating that interference is position specific.

### 4.3 Quantitative model predictions of asymptotic learning

Next, we investigated the accuracy with which asymptotic learning in our empirical data was quantitatively predicted by model simulations. We limited the test of model predictions to the asymptotic part of the learning curves, as early learning likely involves higher-level mechanisms that are not included in the model. As interfering effects of Task A on Task B are absent or limited, model predictions focused on interference by Task B on Task A. We evaluated the agreement between simulated and empirical asymptotic learning curves (sessions 8-15) in the baseline and test conditions for each of our experiments using residual mean-squared error (RMSE).

We first show results from Experiment 1, in which no interference was expected. The model predicted that test thresholds (Task A_T_, solid light-green line figure 5a) would decrease below baseline thresholds (Task A_B_, dashed dark-green line figure 5a). This fits with the expected absenceof interferencefrom Task B, and expected continuation of learning in Task A_T_ from the level achieved in baseline measurements in Task A_B_. The mean difference of test minus baseline predicted by the model was - 0.185 ln-units (figure 5c, dark-green dotted line). This prediction was close to the empirical data (figure 5c, light-green dotted line) with a corresponding empirical mean difference of - 0.226 ln-units (95 % CI [−0.495 0.042]). The close correspondence of empirical and simulated data is also evident in a direct comparison of simulated and empirical results in Task A, which agreed well for baseline (RMSE = 0.056, dashed lines in figures 5a,b) and test (RMSE = 0.071, solid lines figures 5a,b). For comparison, the RMSE between empirical baseline learning curves in Experiments 1 and 2 was 0.115 (compare dashed lines between figures 4a and 4d), thus providing a ballpark estimate for good agreement. A one-sided t-test (*t*_(7)_ = −1.65, *p* = 0.07) marginally failed to show that the empirical mean difference A_T_ – A_B_ was significantly below zero (see figure 5b and light-green dotted line figure 5c).

**Figure 5:**
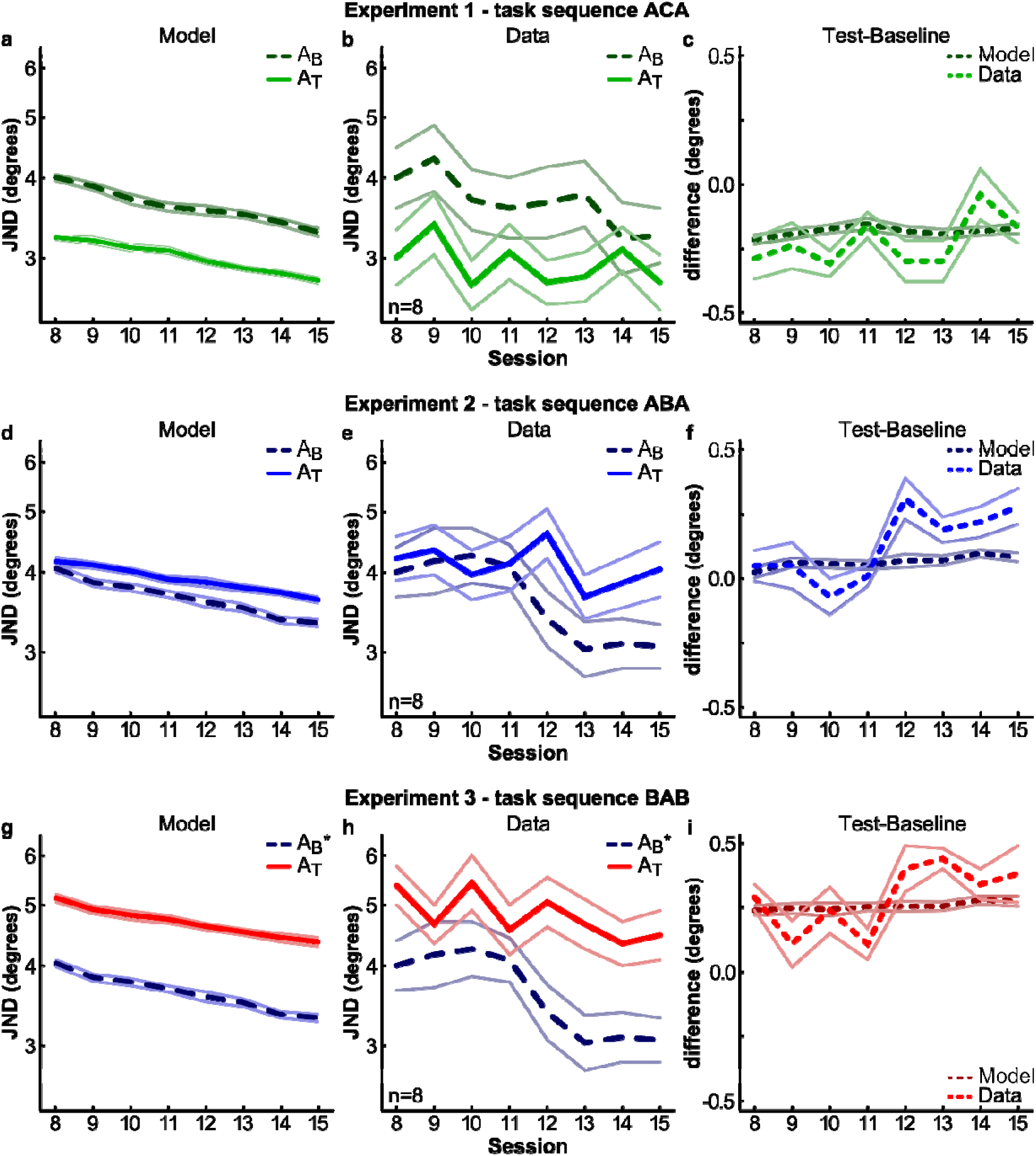
Baseline and test learning curves for asymptotic learning. Model predictions (left column), observed learning asymptotes (middle column) and ln(JND) difference curves for Test-Baseline (right column). Baseline and test learning curves for Task A (135° reference) are compared in Experiments 1 (panels **a-c**), Experiment 2 (panels **d-f**), and Experiment 3 (panels **g-i**). Empirical baselines in **e** and **h** are the same (taken from Experiment 2). **a-c**, Predicted (panel **a**) and observed (panel **b**) asymptotes for Task A_B_ (dashed line) and A_T_ (solid line) in Experiment 1, as well as a direct Test minus Baseline comparison for model (dark dash) and empirical data (light dash; panel **c**). **d-f**, Empirical data versus model comparison for Experiment 2, conventions as in **a-c**. **g-i**, Empirical data versus model comparison for Experiment 3, conventions as in **a-c**. *Please note that the model baseline learning curves in **g** and **d** as well as the empirical baseline learning curves in **h** and **e** are identical (taken from Experiment 2, and therefore shown in blue). For model predictions, the shaded areas represent SEM (the model was trained 25 times in each experiment, see section 2.2 for details); for empirical data, the shaded area represents error (SEM/2). Error values are purely for illustration as all statistical tests were within-subjects.

The model predicted for Experiment 2 (sequence ABA) that overall, asymptotic ln-transformed thresholds in Task A_T_ would be highly similar (though marginally larger) than those observed in Task A_B_(compare solid and dashed blue lines in figure 5d). Accordingly, the empirical asymptotes for Tasks A_T_ and A_B_ (figure 5e) were significantly equivalent (*t*_1(7)_ = −6.595, *t*_2(7)_ = 4.260, *p =* 0.002). The predicted mean difference (Δ_model_ = 0.065, dotted dark-blue line figure 5f) was statistically matched to the empirical mean difference (Δ_data_ = 0.130, dotted light-blue dotted line figure 5f), and fell within the 95% confidence interval of the empirical mean difference [−0.074 0.334]. A direct comparison of the empirical and simulated results for Task A_B_ also showed good agreement with RMSE = 0.095 (compare dashed dark-blue lines in figure 5d,e), and the same held for Task A_T_ with RMSE = 0.087 (compare solid bright-blue lines in figures 5d,e). Overall, predictions and data showed good agreement. Note that the model prediction of largely equivalent asymptotes does not contrast with our earlier comparison of performance in the last three sessions. The model also predicted that asymptotic thresholds in Task A_T_ in Experiment 2 (solid blue line in figure 5d) would eventually diverge from asymptotic thresholds in Task A_B_ (dashed blue line in figure 5d), which was confirmed by the psychophysical data (compare solid and dashed lines in figure 5e, details in Section 4.2).

For Experiment 3 (BAB sequence), the model (see figure 5g) predicted a large increase of test thresholds in Task A_T_(solid red line) compared to baseline thresholds in Task A_B_ (dashed dark-blue line, taken from Experiment 2, justification in legend figure 5). This was confirmed by the empirical data (figure 5h). The predicted mean difference Δ_model_ = 0.255 closely matched the empirical mean difference Δ_data_ = 0.287 (figure 5i). This predicted mean difference (figure 5i, dotted dark-red line) fell within the confidence interval for the empirical mean difference (figure 5i, dotted light-red line) (95 % CI [0.045 0.530]). A one-sided t-test revealed that the mean A_T_ – A_B_ difference between empirical asymptotes was significantly larger than zero (*t*_(7)_ = 2.32, *p* = 0.027; figure 5h,i). A direct comparison of empirical and simulated data for the test condition (Task A_T_, solid red lines figures 4g,h) showed excellent agreement (RMSE = 0.061), and the same was true for the baseline data (identical to those used in Experiment 2, figures 4d,e).

### 4.4 Training balance predicts performance levels

The model predicted that asymptotic performance at the 135° reference orientation during baseline (Task A_B_) or test (Task A_T_) should depend on the balance of training quantity between Task A (135°) and B (105° & 165°). Figure 6a shows that the lowest thresholds were expected (grey bars) and observed (black bars) for Task A_T_ in Experiment 1 (A_T_ | E1). This is because in Experiment 1, there was no interference in the second training period, such that the asymptotic performance at the 135° reference orientation resulted from 30 sessions of training, countered by 0 sessions of interference (30:0 data point in figure 6b). Slightly higher thresholds were predicted and observed in Task A_B_ in Experiments 1 and 3 (A_B_ | E1 and A_B_ | E3 in figure 6a), where the asymptote resulted from 15 training sessions not countered by any interference (15:0 in figure 6b). A further threshold increase was predicted and observed for Task A_T_ in Experiment 2 (A_T_ | E2 in figure 6a), because in this case asymptotic performance resulted from 30 training sessions in Task A, countered however by 15 interfering training sessions in Task B (30:15 in figure 6b). The highest predicted and empirical thresholds were encountered for task A_T_in Experiment 3 (A_T_ | E3 in figure 6a), where 15 sessions in Task B preceded 15 session in Task A_T_ (15:15 in figure 6b). In figure 6a, all predicted values fell within the 95 % confidence intervals of the corresponding empirical data. In figure 6b, the model predicted a log-linear relationship between asymptotic threshold levels and the balance of training between Tasks A and B, with a predicted slope β_model_of 0.10. This was almost identical to the empirically observed slope (β_data_ = 0.11, 95 % CI [0.05 0.17]). A repeated measures linear trend analysis on the ln-transformed empirical data also confirmed the log-linear trend in our data (*F*_(1,7)_ = 11.608, *p* = .011). These results affirm the view that the most extensively trained skills will dominate tuning curve shapes, at the cost of other, less trained skills.

**Figure 6:**
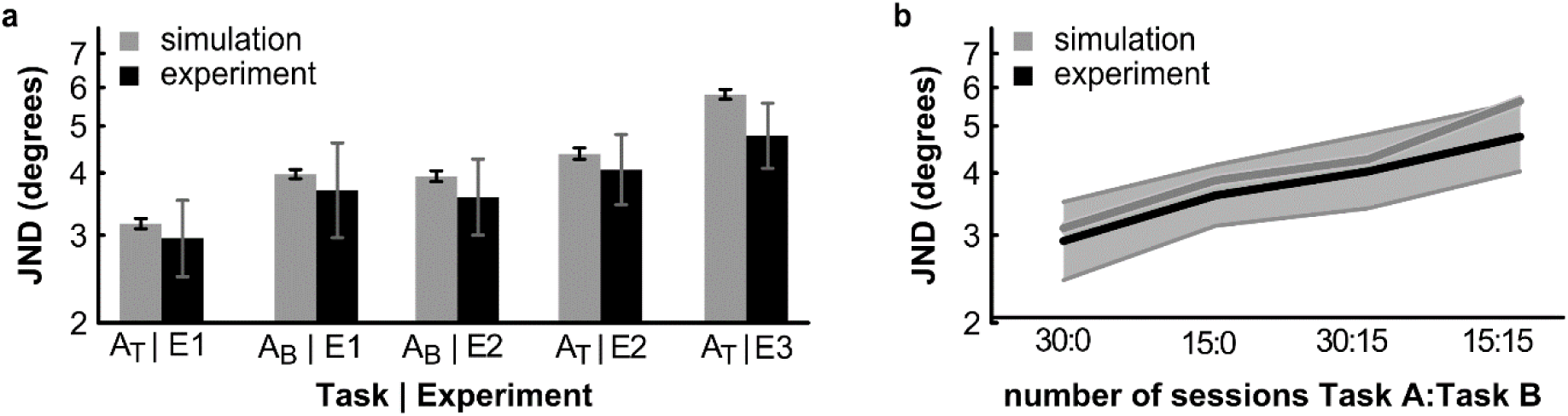
Comparison of performance in different experiments, highlighting the role of the relative amount of training on each task. Task A asymptotes are obtained at reference 135°, during baseline (Task A_B_) or test (Task A_T_) periods. Task B corresponds to training at reference of 105° and 165°. **a**, Mean threshold levels for asymptotic learning (session 8-15) at a 135° reference orientation in Task A_B_ and Task A_T_ of Experiment 1 (E1), as well as Task A_B_and Task A_T_ of Experiment 2 (E2), and Task A_T_ of Experiment 3 (E3; see figure2 in section 3.2 for task conditions). **b**, The balance in number of sessions performed in Task A vs B determines thresholds in Task A. The more the balance is in favor of Task A, the lower the asymptotic thresholds are in Task A. The log-linear trend predicted by the model (grey plot) was similar to that present in the empirical data (black line, grey region shows SEM). Note that the 15:0 balance is equally exhibited by Task A_B_ in E1 and E2 and we only included data of E1 in our analyses to keep sample sizes consistent (choosing Task A_B_ of E2 instead does not alter the effect).

## 5 Discussion

Our data indicate that highly trained skills have no intrinsic longevity and can be erased by training another, sufficiently similar skill. Our psychophysical data on orientation discrimination learning and previously observed tuning curve changes (Raiguel et al., 2006; Schoups et al., 2001) can be well understood by a computational model in which learning of an initial task is due to enhanced inhibition within a part of a neural network and in which interference (resulting from training in a slightly different task) is due to added inhibition in an adjacent part of the network. Hence, interference was created without erasing the connections left behind by original learning. In addition, in line with the incremental nature of the Hebbian learning rule used in our model, the balance of inhibition among parts in the network recruited by different tasks was expected to be related to the difference in trials devoted to each task. This idea was supported by our empirical finding that the efficacy of behavioral interference depended on the difference in the number of sessions devoted to the originally trained task and to the interfering task. Hence, even expert skills are vulnerable to interference, the magnitude of which depends on the extent of training on the similar, interfering skill. Remarkably, the data in our paradigm were excellently predicted by a computational model in which interference occurred without erasing existing connectivity from the originally learned expert skill.

This is the first study that tested interference among skills trained sequentially in a considerable number of sessions. In several previous studies, interference was investigated among two skills trained in a single day. These studies have reported widely ranging time intervals over which interference occurs, from 1-4h (Brashers-Krug et al., 1996; Shadmehr & Brashers-Krug, 1997; J.-Y. Zhang et al., 2008) up to several days (Caithness et al., 2004; Goedert & Willingham, 2002; Yotsumoto, Watanabe, Chang, & Sasaki, 2013). The short-lasting time window following training amenable for eliciting behavioral interference fits roughly with the expression window of late genes controlling synaptic plasticity (Caroni, Chowdhury, & Lahr, 2014; Igaz, Bekinschtein, Vianna, Izquierdo, & Medina, 2004). This can be seen as support for the idea that time-limited latent consolidation is the basis for behavioral interference. By contrast, studies reporting interference over days (Caithness et al., 2004; Goedert & Willingham, 2002; Yotsumoto et al., 2013) have been used to support a view of consolidation in which modified connections remain malleable at all times. This idea has been proposed both in the domain of hippocampal learning (Forcato et al., 2007; Hupbach, Gomez, & Nadel, 2009; Lewis, 1979) and skill learning (Walker, Brakefield, Hobson, & Stickgold, 2003) and holds that memory traces can at any time become modified when reactivated through new experiences. The flexible modulation of neural connectivity through spine dynamics (Hofer, Mrsic-Flogel, Bonhoeffer, & Hübener, 2006; Smith, Heynen, & Bear, 2009) is certainly in support of the modifiability of memory traces. The difficulty of pattern separation in cortex (McClelland et al., 1995; O’Reilly & Rudy, 2001) combined with the malleability of synaptic connectivity could thus form the basis for interference.

In addition to interference with ongoing consolidation, and interference due to the malleability of consolidated synapses, behavioral interference could also rely on a third mechanism. We suggest this mechanism exploits the balance of inhibition within lateral networks that influence the neuronal tuning to relevant values on a particular task-relevant dimension (here, orientation). Because orientation tuning is to a significant extent derived from interactions of any given neuron with its neighbors (Shapley et al., 2003; Sillito, 1975), changes of inhibitory connections in a subpart of the network can produce asymmetric tuning curves that enhance psychophysical performance. Changes in another subpart of the network (due to another task) can restore the balance of inhibitory inputs, thus interfering with prior learning. Thus, for a given tuning curve, two sufficiently similar tasks would pull it in opposite directions (see figure 1). These opposite effects are achieved by strengthening the lateral inhibition in separate subsets of connections among the neighbors influencing a given neuron’s tuning curve, rather than by erasing inhibitory connections already set by the first learning experience. This mechanism may be most strongly recruited in low-level visual areas during and after asymptotic skill learning, as suggested by tuning changes following prolonged training in orientation discrimination (Raiguel et al., 2006; Schoups et al., 2001; Yang & Maunsell, 2004).

The mechanism of balancing network inhibition provides a new perspective on why some studies showed behavioral interference over long time intervals of a day or more (Been et al., 2011; Caithness et al., 2004; Goedert & Willingham, 2002; Yotsumoto et al., 2013), rather than the short intervals of a few hours reported in other studies (Brashers-Krug et al., 1996; Shadmehr & Brashers-Krug, 1997; J.-Y. Zhang et al., 2008). For example, Been et al. (2011) alternated daily training between two orientation discrimination tasks (which we here referred to as Task A and B) and found the same interference for delays between tasks from 0h up to 24h. This is not in line with the notion of time-limited consolidation, but could be due to the fact that performance in the two tasks relies on the modification of different subsets of inhibitory connections in the contributing neural network. Likewise, Caithness et al.’s (2004) finding of interference among different visuo-motor tasks trained in sessions separated by up to a week, may be due to the balancing out of inhibition in a task-relevant network of direction-tuned motor neurons (Georgopoulos, Schwartz, & Kettner, 1986). The same explanatory model may also help to understand data from other studies demonstrating interference over long time intervals (Goedert & Willingham, 2002; Yotsumoto et al., 2013).

How exactly the balancing of network inhibition is achieved in the cortex remains an open question. In the light of evidence that inhibition shapes orientation tuning (Sillito, 1975), and that inhibitory interneurons in V1 become increasingly selective with learning (Khan et al., 2018), we modeled (re-)balancing as an increase in inhibitory connections. We could have achieved the same effect if instead we had chosen to decrease excitatory connections (cf. Teich & Qian, 2003). Our model consists of a set of rate neurons forming both excitatory and inhibitory connections with each other. This is a considerable simplification because neurons are distinctly excitatory or inhibitory, as expressed in Dale’s law (Eccles, 1976). In a more realistic network model, balancing of network inhibition might be achieved by a) reducing synaptic coupling among excitatory neurons, b) increasing synaptic coupling from excitatory to inhibitory neurons, c) increasing synaptic coupling from inhibitory to excitatory neurons, or d) reducing synaptic coupling among inhibitory neurons. We implicitly assumed either b) or c) to be the case. However, which of these mechanisms regulates network inhibition is a question that requires further research.

A related question pertains to how learning could occur in response to negative as opposed to positive feedback; i.e. after incorrect as opposed to correct trials. The latter is typically assumed to result from reward-related neuromodulators such as dopamine gating plasticity (Glimcher, 2011). The former would require a punishment-related neuromodulator. A candidate neurotransmitter potentially able to fulfill this role is serotonin whose levels are elevated in response to cues reflective of punishment and aversive stimuli (Cools, Nakamura, & Daw, 2011; Crockett, Clark, & Robbins, 2009; Evers et al., 2005). Vasoactive intestinal peptide expressing (VIP+) interneurons in early visual cortex express the serotonin receptor 5HT3aR (van Versendaal & Levelt, 2016) and may thus be activated by serotonin released in response to incorrect trials. Given that VIP+ interneurons inhibit somatostatin expressing (SOM) interneurons, which in turn inhibit pyramidal cells as well as parvalbumin expressing (PV+) interneurons (Xu, Jeong, Tremblay, & Rudy, 2013), disinhibition of pyramidal cells or PV+ interneurons in response to increased VIP+ neuron activity has been suggested to modulate experience-dependent plasticity in early visual cortex (van Versendaal & Levelt, 2016). Balancing of network inhibition either according to mechanisms b) or c) could thus potentially be modulated by serotonin. We do not wish to claim that this particular mechanism explains how error-triggered Hebbian learning is implemented in visual cortex. Rather, we only wish to highlight that our modeling choice is biologically plausible as this or a comparable mechanism is principally conceivable given current evidence.

Retinotopic areas other than V1 may show training-induced orientation tuning changes (Raiguel et al., 2006; Yang & Maunsell, 2004). Hence, V1 is not the sole contributor to the observed asymptotic learning and interference (as also recognized in Schoups et al., 2001). The modelled tuning changes during asymptotic learning in our V1-inspired model may thus have been larger than what we might have seen if other areas had been included in the model. Nevertheless, the ability of our relatively simple model to accurately predict major aspects of learning and behavioral interference indicates that our observations to a significant extent are mediated by orientation tuning changes due to altered recurrent interactions in mid-to-low levelsofthe visual system.

As an alternative to altering recurrent interactions within low-level visual areas during asymptotic learning, learning by altering feedforward connections may also lead to behavioral interference. However, the expected interference from such a feedforward learning model likely differs not only quantitatively but also qualitatively from what we observed here. Models relying on feedforward mechanisms to sharpen tuning curves have been shown to fail to explain the shift in the peak of orientation tuning curves towards the reference orientation (Teich & Qian, 2010). Given that these shifts account, at least in part, for flattening of tuning curve flanks facing away from the reference orientation, they are likely to shape interference effects. Furthermore, if the same feedforward weights need to be sharpened to increase sensitivity around similar orientations, it is likely that extensive initial training at one set of orientations will interfere with subsequent learning using other orientations. In this scenario, there would be strong interference irrespective of the order of Tasks A and B. In contrast, the recurrent mechanism employed here does not predict such generalized proactive interference, and it is indeed also absent from our psychophysical data (in figure 4 compare panels d and e, and panels g and h). This argument extends to read-out effects, which are similarly likely to produce proactive interference if discrimination around different references needs to rely on a single, shared set of feedforward weights. In a more complicated read-out scenario with multiple read-out units that do not share feedforward weights, no interference at all is expected, which also conflicts with our empirical data. Notably, it has recently been shown that exposing a purely feedforward convolutional neural network to orientation discrimination training can sharpen tuning curves in early layers of the model (reflecting V1- and V2-like processing; Wenliang & Seitz, 2018). The convolutional neural network has, however, not been used to investigate interference. Given that it uses a large number of independent filters at all hierarchical levels of the network, we suspect it would not exhibit the qualitative interference effects observed in our study. Thus, while read-out effects likely would have contributed to learning in our study (Hochstein & Ahissar, 2002; Law & Gold, 2008), they cannot easily account for the interference effects we observed in the later, asymptotic parts of the learning curves. These interference effects are parsimoniously explained by assuming recurrent interactions among orientation-tuned neurons (Teich & Qian, 2003) restricted to mid-to-low levels in the visual system (A Karni & Bertini, 1997; Avi Karni & Sagi, 1993; Schoups et al., 2001), and compatible with views that permit learning-induced changes in both lower and higher levels of the visual system (Dosher, Jeter, Liu, & Lu, 2013; Hochstein & Ahissar, 2002). Nevertheless, future efforts to model the observed behavioral interference should combine learning-induced plasticity in low-level sensory cortex with a plastic read-out mechanism.

The psychophysical data showed that following training in Task A, intervening training in Task B led to a substantial threshold increase in the first sessions of returning to Task A (figure 4, d-f). It is possible that the elevated thresholds indicate a ‘reset’ in which performance depends again on mid to high-level visual areas that can support coarse orientation discrimination (Hochstein & Ahissar, 2002). Alternatively, it is possible that upon returning to Task A, information continues to be read out from the lowest level, and that the quasi-naïve network state at low levels (figure 3c) is the reason why thresholds are high immediately after returning to Task A. This is another open question that should be investigated in models that combine low-level and read-out plasticity.

The behavioral interference effects in our study during asymptotic learning were specific to the manipulations imposed in different visual field quadrants (compare asymptotes in figure 4c and 4f). This appears to conflict with studies showing substantial or complete generalization of visual skill learning (e.g., Xiao et al., 2008; J.-Y. Zhang et al., 2010; T. Zhang, Xiao, Klein, & Levi, 2010). However, experiments in orientation discrimination that showed generalization among peripheral locations were limited to early learning (T. Zhang et al., 2010), and other experiments indicated that generalization among peripheral locations does not include progress made during asymptotic learning (Lange & De Weerd, 2018). In addition, studies showing strong generalization (e.g., Xiao et al., 2008; J.-Y. Zhang et al., 2010; T. Zhang, Xiao, Klein, & Levi, 2010) often limited the length of learning curves, or used particular training procedures that we did not apply in the present study (for a full discussion, see Lange & De Weerd, 2018). At present, there is no evidence indicating that the asymptotic performance enhancements in expert orientation discrimination acquired after prolonged learning (15 sessions or more) show substantial generalization among peripheral visual field locations. Hence, we suggest there is no conflict between the position specific interference during late learning in our study and the findings of generalization in other studies using other paradigms (e.g., Xiao et al., 2008; J.-Y. Zhang et al., 2010; T. Zhang, Xiao, Klein, & Levi, 2010). Asymptotic learning, rather than being amenable to substantial generalization, has been demonstrated to induce localized plasticity in low-level visual areas (Raiguel et al., 2006; Schoups et al., 2001; Yang & Maunsell, 2004), as well as skill that is specific for visual field position (Lange & De Weerd, 2018; Lange, Lowet, Roberts, & De Weerd, 2018; Schoups et al., 1995, 2001). The specificity of interference effects on asymptotic learning of orientation discrimination is in accordance with the specificity of asymptotic learning itself.

We also found position specificity of the interference effects in the early part of the learning curves of Experiment 1 (compare sessions 31-33 in figure 4c and 4f). To the extent that the lower-level network immediately following the return to Task A (figure 4f, first sessions) is close to its naïve state (figure 3c), this position specific interference effect may simply reflect the accompanying reduction in informativeness of tuning curves and hence rely on low-level visual cortex. It is possible, however, that this observation instead reveals a contribution to position specificity from read-out networks extending beyond low levels of the visual system. Earlier work has shown that during early perceptual learning, low-level visual areas may already be included in a larger network supporting skill acquisition and memory formation (De Weerd et al., 2012; Schwartz, Maquet, & Frith, 2002; Yotsumoto, Sasaki, et al., 2009; Sarabi et al., 2018). It is possible that when the context of the training makes a given position relevant, read-out itself may develop position specificity. This view is in line with the general idea that the read-out mechanisms in perceptual learning are flexible and rule-based (Xiao et al., 2008; J.-Y. Zhang et al., 2010; T. Zhang et al., 2010), and can be shaped to optimize read-out in a single or multiple locations depending on task requirements (Eckstein, Abbey, Pham, & Shimozaki, 2004). Hence, it is possible that different magnitudes of interference in different visual field locations are embedded at least partly in specific cortico-cortical connectivity that links the task-relevant retinotopic regions in low-level visual cortex with one (or possibly more) read-out mechanisms. Further research is required to test the contributions of higher and lower levels of the visual system during different learning phases to position specificity of acquisition, performance and interference.

The present study has several limitations. It did not include a read-out mechanism, which in future work should be added to the recurrent mechanism modeled in a prototypical lower-level visual area. Future modeling should also entail a more biologically plausible implementation of the decision process. This may be modeled by a winner-take-all attractor neural network as put forth by Wang (2002), potentially supplemented by a criterion (c.f. Insabato, Pannunzi, Rolls, & Deco, 2010) reflecting the internal reference. In the present study, orientation discrimination decisions were performed using signal detection theory (Green & Swets, 1966), which constitutes an approach to the behavioral manifestation of the decision, but not of the decision process itself.

Despite limitations, our empirical and model data yielded several insights. First, our data suggest that highly trained skills are not more stable than less trained skills. Rather, the stability of highly trained skills is conditional on the lack of interfering expertise. Specifically, highly trained skills may remain stable simply because in most domains of skill learning it is unlikely that new skills would be trained that are similar enough and trained for long enough to interfere with established expertise. However, when a new skill is trained that accesses a neuronal population strongly overlapping with the neuronal population where the older expertise is stored, the older expert skill will become damaged to the extent that more training is devoted to the new skill. Second, our results also highlight the importance of separately considering perceptual skill in terms of altered neuronal tuning, and in terms of the altered network connectivity that corresponds to the ‘memory trace’, as the insights from one perspective do not necessarily translate directly to the other perspective. In the present study, a V1 model accurately predicting interference between two orientation discrimination tasks indicates that although interference can undo previously established changes of neuronal tuning curves, this does not necessarily translate into erasure of previously established neuronal connectivity. Third, our empirical and modeling work further indicates that memory traces understood as altered synaptic interactions do not exert their influence in isolation but only in the context of other memory traces. This is because neuronal tuning curves sample these changes in connectivity induced by new learning in the context of other connectivity that may reflect older learning experiences. Tuning curves hence form the interface between discrimination behavior and connectivity patterns representing the perceptual skill. Therefore, further investigations of the precise relationships between local network connectivity changes and neuronal tuning may increase our understanding of the mechanisms of skill learning as well as the way in which skills can interfere with each other.

## Acknowledgements

The authors would like to thank Joel Reithler, Michelle Moerel and Vaishnavi Narayanan for their helpful comments. Author GL was supported by an NWO grant (022.001.036) to FPN Graduate School for Cognitive and Clinical Neuroscience. Author MS was supported by the European Union’s Horizon 2020 research and innovation programme under grant agreement n. 720270 (HBP SGA1) and 737691 (HBP SGA2) and by the European Research Council under the European Union’s Seventh Framework Programme (<u>ERC- 2010-AdG</u>, ERC grant agreement no. <u>269853</u>). Author PDW was supported by an NWO VICI grant (453.04.002).

## References

Bahrick, H. P. (1984). Semantic memory content in permastore: Fifty years of memory for Spanish learned in school. Journal of Experimental Psychology: General, 113(1), 1–29. https://doi.org/10.1037/0096-3445.113.1.1

Been, M., Jans, B., & De Weerd, P. (2011). Time-limited consolidation and task interference: no direct link. The Journal of Neuroscience□: The Official Journal of the Society for Neuroscience, 31(42), 14944–14951. https://doi.org/10.1523/JNEUROSCI.1046-11.2011

Brashers-Krug, T., Shadmehr, R., & Bizzi, E. (1996). Consolidation in human motor memory. Nature, 382(6588), 252–255.

Caithness, G., Osu, R., Bays, P., Chase, H., Klassen, J., Kawato, M., … Flanagan, J. R. (2004). Failure to consolidate the consolidation theory of learning for sensorimotor adaptation tasks. The Journal of Neuroscience, 24(40), 8662–8671.

Caroni, P., Chowdhury, A., & Lahr, M. (2014). Synapse rearrangements upon learning: from divergent–sparse connectivity to dedicated sub-circuits. Trends in Neurosciences, 37(10), 604–614.

Cools, R., Nakamura, K., & Daw, N. D. (2011). Serotonin and Dopamine: Unifying Affective, Activational, and Decision Functions. Neuropsychopharmacology, 36(1), 98–113. https://doi.org/10.1038/npp.2010.121

Crockett, M. J., Clark, L., & Robbins, T. W. (2009). Reconciling the role of serotonin in behavioral inhibition and aversion: acute tryptophan depletion abolishes punishment-induced inhibition in humans. The Journal of Neuroscience□: The Official Journal of the Society for Neuroscience, 29(38), 11993–9. https://doi.org/10.1523/JNEUROSCI.2513-09.2009

Daniel, P. M., & Whitteridge, D. (1961). The representation of the visual field on the cerebral cortex in monkeys. The Journal of Physiology, 159(2), 203–221.

De Weerd, P., Peralta, M. R., Desimone, R., & Ungerleider, L. G. (1999). Loss of attentional stimulus selection after extrastriate cortical lesions in macaques. Nature Neuroscience, 2(8), 753–758.

Dosher, B. A., Jeter, P., Liu, J., & Lu, Z.-L. (2013). An integrated reweighting theory of perceptual learning. Proceedings of the National Academy of Sciences, 110(33), 13678–13683. https://doi.org/10.1073/pnas.1312552110

Dougherty, K. M., & Johnston, J. M. (1996). Overlearning, Fluency, and Automaticity. The Behavior Analyst, 19(2), 289–292. https://doi.org/10.1007/BF03393171

Dragoi, V., Sharma, J., & Sur, M. (2000). Adaptation-Induced Plasticity of Orientation Tuning in Adult Visual Cortex. Neuron, 28(1), 287–298. https://doi.org/10.1016/S0896-6273(00)00103-3

Eccles, J. (1976). From electrical to chemical transmission in the central nervous system. Notes and Records of the Royal Society of London, 30(2), 219–230. https://doi.org/10.1098/rsnr.1976.0015

Eckstein, M. P., Abbey, C. K., Pham, B. T., & Shimozaki, S. S. (2004). Perceptual learning through optimization of attentional weighting: Human versus optimal Bayesian learner. Journal of Vision, 4(12), 3. https://doi.org/10.1167/4.12.3

Ericsson, K. A., & Lehmann, A. C. (1996). Expert and Exceptional Performance: Evidence of Maximal Adaptation to Task Constraints. Annual Review of Psychology, 47(1), 273–305. https://doi.org/10.1146/annurev.psych.47.1.273

Evers, E. A. T., Cools, R., Clark, L., van der Veen, F. M., Jolles, J., Sahakian, B. J., & Robbins, T. W. (2005). Serotonergic Modulation of Prefrontal Cortex during Negative Feedback in Probabilistic Reversal Learning. Neuropsychopharmacology, 30(6), 1138–1147. https://doi.org/10.1038/sj.npp.1300663

Farr, M. J. (1987). The Long-Term Retention of Knowledge and Skills: A Cognitive and Instructional Perspective. https://doi.org/10.1007/978-1-4612-1062-7

Ferster, D., Chung, S., & Wheat, H. (1996). Orientation selectivity of thalamic input to simple cells of cat visual cortex. Nature, 380(6571), 249–252. https://doi.org/10.1038/380249a0

Fisk, A. D., Hertzog, C., Lee, M. D., Rogers, W. A., & Anderson-Garlach, M. (1994). Long-term retention of skilled visual search: Do young adults retain more than old adults? Psychology and Aging, 9(2), 206–215. https://doi.org/10.1037/0882-7974.9.2.206

Forcato, C., Burgos, V. L., Argibay, P. F., Molina, V. A., Pedreira, M. E., & Maldonado, H. (2007). Reconsolidation of declarative memory in humans. Learning & Memory, 14(4), 295–303.

Gattass, R., Sousa, A. P. B., & Rosa, M. G. P. (1987). Visual topography of V1 in the Cebus monkey. Journal of Comparative Neurology, 259(4), 529–548.

Georgopoulos, A. P., Schwartz, A. B., & Kettner, R. E. (1986). Neuronal population coding of movement direction. Science (New York, N.Y.), 233(4771), 1416–9. Retrieved from http://www.ncbi.nlm.nih.gov/pubmed/3749885

Glimcher, P. W. (2011). Understanding dopamine and reinforcement learning: The dopamine reward prediction error hypothesis. Proceedings of the National Academy of Sciences, 108(Supplement 3), 15647–15654. https://doi.org/10.1073/PNAS.1014269108

Goedert, K. M., & Willingham, D. B. (2002). Patterns of interference in sequence learning and prism adaptation inconsistent with the consolidation hypothesis. Learning & Memory, 9(5), 279–292.

Green, D. M., & Swets, J. A. (1966). Signal detection theory and psychophysics. New York: John Wiley, Inc.

Hagman, J. D., & Rose, A. M. (1983). Retention of Military Tasks: A Review. Human Factors: The Journal of the Human Factors and Ergonomics Society, 25(2), 199–213. https://doi.org/10.1177/001872088302500207

Healy, A. F., Clawson, D. M., McNamara, D. S., Marmie, W. R., Schneider, V. I., Rickard, T. C., … Bourne, L. E. (1993). The Long-Term Retention of Knowledge and Skills. In Psychology of Learning and Motivation - Advances in Research and Theory (Vol. 30, pp. 135–164). Academic Press Inc. https://doi.org/10.1016/S0079-7421(08)60296-0

Healy, A. F., Fendrich, D. W., & Proctor, J. D. (1990). Acquisition and retention of a letter-detection skill. *Journal of Experimental Psychology: Learning*, Memory, and Cognition, 16(2), 270–281. https://doi.org/10.1037/0278-7393.16.2.270

Hikosaka, O., Rand, M. K., Nakamura, K., Miyachi, S., Kitaguchi, K., Sakai, K., … Shimo, Y. (2002). Long-term retention of motor skill in macaque monkeys and humans. Experimental Brain Research, 147(4), 494–504. https://doi.org/10.1007/s00221-002-1258-7

Hochstein, S., & Ahissar, M. (2002). View from the top: Hierarchies and reverse hierarchies in the visual system. Neuron, 36(5), 791–804.

Hofer, S. B., Mrsic-Flogel, T. D., Bonhoeffer, T., & Hübener, M. (2006). Lifelong learning: ocular dominance plasticity in mouse visual cortex. Current Opinion in Neurobiology, 16(4), 451–459. https://doi.org/10.1016/j.conb.2006.06.007

Hubel, D. H., & Wiesel, T. N. (1959). Receptive fields of single neurones in the cat’s striate cortex. The Journal of Physiology, 148(3), 574–591. Retrieved from http://onlinelibrary.wiley.com/doi/10.1113/jphysiol.1959.sp006308/full

Hupbach, A., Gomez, R., & Nadel, L. (2009). Episodic memory reconsolidation: updating or source confusion? Memory, 17(5), 502–510.

Igaz, L. M., Bekinschtein, P., Vianna, M. M. R., Izquierdo, I., & Medina, J. H. (2004). Gene expression during memory formation. Neurotoxicity Research, 6(3), 189–203.

Insabato, A., Pannunzi, M., Rolls, E. T., & Deco, G. (2010). Confidence-Related Decision Making. Journal of Neurophysiology, 104(1), 539.

Karni, A., & Bertini, G. (1997). Learning perceptual skills: behavioral probes into adult cortical plasticity. Current Opinion in Neurobiology, 7(4), 530–535. Retrieved from http://www.sciencedirect.com/science/article/pii/S0959438897800335

Karni, A., & Sagi, D. (1993). The time course of learning a visual skill. Nature, 365(6443), 250–252. https://doi.org/10.1038/365250a0

Kaufman, S. B., & Kaufman, J. C. (2007). Ten Years to Expertise, Many More to Greatness: An Investigation of Modern Writers. The Journal of Creative Behavior, 41(2), 114–124. https://doi.org/10.1002/j.2162-6057.2007.tb01284.x

Khan, A. G., Poort, J., Chadwick, A., Blot, A., Sahani, M., Mrsic-Flogel, T. D., & Hofer, S. B. (2018). Distinct learning-induced changes in stimulus selectivity and interactions of GABAergic interneuron classes in visual cortex. Nature Neuroscience, 1. https://doi.org/10.1038/s41593-018-0143-z

King, P. D., Zylberberg, J., & DeWeese, M. R. (2013). Inhibitory Interneurons Decorrelate Excitatory Cells to Drive Sparse Code Formation in a Spiking Model of V1. Journal of Neuroscience, 33(13), 5475–5485. https://doi.org/10.1523/JNEUROSCI.4188-12.2013

Koch, G., Ponzo, V., Lorenzo, F. Di, Caltagirone, C., & Veniero, D. (2013). Hebbian and Anti-Hebbian Spike-Timing-Dependent Plasticity of Human Cortico-Cortical Connections. The Journal of Neuroscience, 33(23), 9725–9733. https://doi.org/10.1523/JNEUROSCI.4988-12.2013

Krakauer, J. W., Ghilardi, M.-F., & Ghez, C. (1999). Independent learning of internal models for kinematic and dynamic control of reaching. Nature Neuroscience, 2(11), 1026–1031. https://doi.org/10.1038/14826

Lange, G., & De Weerd, P. (2018). Limited transfer of visual skill in orientation discrimination to locations treated by pre-testing and subliminal exposure. Vision Research, 143, 103–116. https://doi.org/10.1016/j.visres.2017.06.018

Lange, G., Lowet, E., Roberts, M. J., & De Weerd, P. (2018). Within-quadrant position and orientation specificity after extensive orientation discrimination learning is related to performance gains during late learning. PLOS ONE, 13(9), e0201520. https://doi.org/10.1371/journal.pone.0201520

Law, C. T., & Gold, J. I. (2008). Neural correlates of perceptual learning in a sensory-motor, but not a sensory, cortical area. Nature Neuroscience, 11(4), 505–513. Retrieved from http://dx.doi.org/10.1038/nn2070

Lewis, D. J. (1979). Psychobiology of active and inactive memory. Psychological Bulletin, 86(5), 1054.

Mara, C. A., & Cribbie, R. A. (2012). Paired-Samples Tests of Equivalence. Communications in Statistics - Simulation and Computation, 41(10), 1928–1943. https://doi.org/10.1080/03610918.2011.626545

McClelland, J. L., McNaughton, B. L., & O’Reilly, R. C. (1995). Why there are complementary learning systems in the hippocampus and neocortex: insights from the successes and failures of connectionist models of learning and memory. Psychological Review, 102(3), 419.

Ni, A. M., & Maunsell, J. H. R. (2010). Microstimulation reveals limits in detecting different signals from a local cortical region. Current Biology□: CB, 20(9), 824–8. https://doi.org/10.1016/j.cub.2010.02.065

O’Reilly, R. C., & Rudy, J. W. (2001). Conjunctive representations in learning and memory: principles of cortical and hippocampal function. Psychological Review, 108(2), 311.

Park, S.-W., Dijkstra, T. M. H., & Sternad, D. (2013). Learning to never forget-time scales and specificity of long-term memory of a motor skill. Frontiers in Computational Neuroscience, 7, 111. https://doi.org/10.3389/fncom.2013.00111

Peres, R., & Hochstein, S. (1994). Modeling perceptual learning with multiple interacting elements: a neural network model describing early visual perceptual learning. Journal of Computational Neuroscience, 1(4), 323–338.

Raiguel, S., Vogels, R., Mysore, S. G., & Orban, G. A. (2006). Learning to See the Difference Specifically Alters the Most Informative V4 Neurons. The Journal of Neuroscience, 26(24), 6589–6602. https://doi.org/10.1523/jneurosci.0457-06.2006

Sarabi, M. T., Aoki, R., Tsumura, K., Keerativittayayut, R., Jimura, K., & Nakahara, K. (2018). Visual perceptual training reconfigures post-task resting-state functional connectivity with a feature-representation region. PLOS ONE, 13(5), e0196866. https://doi.org/10.1371/journal.pone.0196866

Schoups, A. A., Vogels, R., & Orban, G. A. (1995). Human perceptual learning in identifying the oblique orientation: retinotopy, orientation specificity and monocularity. The Journal of Physiology, 483(3), 797–810. https://doi.org/10.1113/jphysiol.1995.sp020623

Schoups, A. A., Vogels, R., Qian, N., & Orban, G. A. (2001). Practising orientation identification improves orientation coding in V1 neurons. Nature, 412(6846), 549–553. Retrieved from http://dx.doi.org/10.1038/35087601

Seaman, M. A., & Serlin, R. C. (1998). Equivalence confidence intervals for two-group comparisons of means. Psychological Methods, 3(4), 403.

Seitz, A. R., Yamagishi, N., Werner, B., Goda, N., Kawato, M., & Watanabe, T. (2005). Task-specific disruption of perceptual learning. Proceedings of the National Academy of Sciences of the United States of America, 102(41), 14895–14900.

Shadlen, M. N., & Newsome, W. T. (1994). Noise, neural codes and cortical organization. Current Opinion in Neurobiology, 4(4), 569–579.

Shadmehr, R., Brandt, J., & Corkin, S. (1998). Time-Dependent Motor Memory Processes in Amnesic Subjects. Journal of Neurophysiology, 80(3), 1590–1597. https://doi.org/10.1152/jn.1998.80.3.1590

Shadmehr, R., & Brashers-Krug, T. (1997). Functional stages in the formation of human long-term motor memory. The Journal of Neuroscience, 17(1), 409–419.

Shadmehr, R., & Holcomb, H. H. (1997). Neural correlates of motor memory consolidation. Science, 277(5327), 821–825.

Shapley, R., Hawken, M., & Ringach, D. L. (2003). Dynamics of orientation selectivity in the primary visual cortex and the importance of cortical inhibition. Neuron, 38(5), 689–99. Retrieved from http://www.ncbi.nlm.nih.gov/pubmed/12797955

Sillito, A. M. (1975). The contribution of inhibitory mechanisms to the receptive field properties of neurones in the striate cortex of the cat. The Journal of Physiology, 250(2), 305–329. https://doi.org/10.1113/jphysiol.1975.sp011056

Simon, H. A., & Chase, W. G. (1973). Programs and probability. American Scientist, 61(4), 394–403. https://doi.org/10.1511/2011.89.106

Smith, G. B., Heynen, A. J., & Bear, M. F. (2009). Bidirectional synaptic mechanisms of ocular dominance plasticity in visual cortex. Philosophical Transactions of the Royal Society of London. Series B, Biological Sciences, 364, 357–367. https://doi.org/10.1098/rstb.2008.0198

Snowden, R. J., Treue, S., & Andersen, R. A. (1992). The response of neurons in areas V1 and MT of the alert rhesus monkey to moving random dot patterns. Experimental Brain Research, 88(2), 389–400.

Softky, W. R., & Koch, C. (1993). The highly irregular firing of cortical cells is inconsistent with temporal integration of random EPSPs. The Journal of Neuroscience, 13(1), 334–350.

Teich, A. F., & Qian, N. (2003). Learning and Adaptation in a Recurrent Model of V1 Orientation Selectivity. Journal of Neurophysiology, 89(4), 2086–2100. https://doi.org/10.1152/jn.00970.2002

Teich, A. F., & Qian, N. (2010). V1 orientation plasticity is explained by broadly tuned feedforward inputs and intracortical sharpening. Visual Neuroscience, 27(1–2), 57–73. https://doi.org/10.1017/S0952523810000039

van Versendaal, D., & Levelt, C. N. (2016). Inhibitory interneurons in visual cortical plasticity. Cellular and Molecular Life Sciences, 73(19), 3677–3691. https://doi.org/10.1007/s00018-016-2264-4

Vogels, R. (1990). Population coding of stimulus orientation by striate cortical cells. Biological Cybernetics, 64(1), 25–31.

Vogels, R., & Orban, G. A. (1985). The effect of practice on the oblique effect in line orientation judgments. Vision Research, 25(11), 1679–1687. https://doi.org/10.1016/0042-6989(85)90140-3

Walker, M. P., Brakefield, T., Hobson, J. A., & Stickgold, R. (2003). Dissociable stages of human memory consolidation and reconsolidation. Nature, 425(6958), 616–620.

Wang, X.-J. (2002). Probabilistic Decision Making by Slow Reverberation in Cortical Circuits. Neuron, 36(5), 955–968. https://doi.org/10.1016/S0896-6273(02)01092-9

Wenliang, L. K., & Seitz, A. R. (2018). Deep Neural Networks for Modeling Visual Perceptual Learning. Journal of Neuroscience, 38(27), 6028–6044. https://doi.org/10.1523/JNEUROSCI.1620-17.2018

Wetherill, G. B., & Levitt, H. (1965). Sequential estimation of points on a psychometric function. British Journal of Mathematical and Statistical Psychology, 18(1), 1–10.

Wickelgreen, W. A. (1972). Trace resistance and the decay of long-term memory. Journal of Mathematical Psychology, 9(4), 418–455.

Xiao, L.-Q., Zhang, J.-Y., Wang, R., Klein, S. A., Levi, D. M., & Yu, C. (2008). Complete Transfer of Perceptual Learning across Retinal Locations Enabled by Double Training. Current Biology, 18(24), 1922–1926. https://doi.org/10.1016/J.CUB.2008.10.030

Xu, H., Jeong, H.-Y., Tremblay, R., & Rudy, B. (2013). Neocortical Somatostatin-Expressing GABAergic Interneurons Disinhibit the Thalamorecipient Layer 4. Neuron, 77(1), 155–167. https://doi.org/10.1016/j.neuron.2012.11.004

Yang, T., & Maunsell, J. H. (2004). The effect of perceptual learning on neuronal responses in monkey visual area V4. The Journal of Neuroscience, 24(7), 1617–1626. https://doi.org/10.1523/JNEUROSCI.4442-03.2004

Yotsumoto, Y., Chang, L., Watanabe, T., & Sasaki, Y. (2009). Interference and feature specificity in visual perceptual learning. Vision Research, 49(21), 2611–2623.

Yotsumoto, Y., Watanabe, T., Chang, L.-H., & Sasaki, Y. (2013). Consolidated learning can be susceptible to gradually-developing interference in prolonged motor learning. Frontiers in Computational Neuroscience, 7. https://doi.org/10.3389/fncom.2013.00069

Zhang, J.-Y., Kuai, S.-G., Xiao, L.-Q., Klein, S. A., Levi, D. M., & Yu, C. (2008). Stimulus Coding Rules for Perceptual Learning. PLoS Biol, 6(8), e197. https://doi.org/10.1371/journal.pbio.0060197

Zhang, J.-Y., Zhang, G.-L., Xiao, L.-Q., Klein, S. A., Levi, D. M., & Yu, C. (2010). Rule-based learning explains visual perceptual learning and its specificity and transfer. The Journal of Neuroscience□: The Official Journal of the Society for Neuroscience, 30(37), 12323–8. https://doi.org/10.1523/JNEUROSCI.0704-10.2010

Zhang, T., Xiao, L.-Q., Klein, S. A., & Levi, D. M. (2010). Decoupling location specificity from perceptual learning of orientation discrimination. Vision Research, 50(4), 368–374. https://doi.org/10.1016/J.VISRES.2009.08.024

